# LY6E protects mice from pathogenic effects of murine coronavirus and SARS-CoV-2

**DOI:** 10.1101/2023.01.25.525551

**Authors:** Katrina B. Mar, Marley C. Van Dyke, Alexandra H. Lopez, Jennifer L. Eitson, Wenchun Fan, Natasha W. Hanners, Bret M. Evers, John M. Shelton, John W. Schoggins

**Affiliations:** Department of Microbiology, University of Texas Southwestern Medical Center, Dallas, TX; Department of Pediatrics, University of Texas Southwestern Medical Center, Dallas, TX; Departments of Pathology and Ophthalmology, University of Texas Southwestern Medical Center, Dallas, TX; Department of Internal Medicine, Histo Pathology Core Division, University of Texas Southwestern Medical Center, Dallas, TX

## Abstract

LY6E is an antiviral protein that inhibits coronavirus entry. Its expression in immune cells allows mice to control murine coronavirus infection. However, it is not known which immune cell subsets mediate this control or whether LY6E protects mice from SARS-CoV-2. In this study, we used tissue-specific Cre recombinase expression to ablate *Ly6e* in distinct immune compartments or in all epiblast-derived cells, and bone marrow chimeras to target Ly6e in a subset of radioresistant cells. Mice lacking *Ly6e* in *Lyz2*-expressing cells and radioresistant *Vav1*-expressing cells were more susceptible to lethal murine coronavirus infection. Mice lacking *Ly6e* globally developed clinical disease when challenged with the Gamma (P.1) variant of SARS-CoV-2. By contrast, wildtype mice and mice lacking type I and type III interferon signaling had no clinical symptoms after SARS-CoV-2 infection. Transcriptomic profiling of lungs from SARS-CoV-2-infected wildtype and *Ly6e* knockout mice revealed a striking reduction of secretory cell-associated genes in infected knockout mice, including *Muc5b*, an airway mucin-encoding gene that may protect against SARS-CoV-2-inflicted respiratory disease. Collectively, our study reveals distinct cellular compartments in which Ly6e confers cell intrinsic antiviral effects, thereby conferring resistance to disease caused by murine coronavirus and SARS-CoV-2.

## Introduction

Host resistance to viral pathogens depends in part on constitutively expressed and infection-induced effector proteins that selectively target distinct steps of the viral replication cycle. In concert with humoral and cellular immune responses, cell intrinsic antiviral effectors limit viral burden and consequently constrain viral pathogenesis. Intracellular viral detection triggers rapid production and extracellular release of type I and III interferons that act in an autocrine and paracrine manner to induce the expression of cytokines, chemokines, and cell-intrinsic effector proteins (Schoggins 2019). While the expression of a subpopulation of effectors proteins are undetectable in the absence of interferons, several proteins categorized as interferon-stimulated genes (ISGs) are prominently expressed without infectious stimuli. These basally expressed cell-intrinsic effectors may serve as an early barrier against viral infection that precedes production of inflammatory cytokines and recruitment of immune cells.

Lymphocyte antigen 6 complex, locus E (LY6E, previously called retinoic acid-inducible gene E [RIG-E], thymic shared antigen 1 [TSA-1], and stem cell antigen 2 [SCA-2]) is constitutively expressed across multiple tissues in humans (Mao et al. 1996) and mice (Bacquin et al. 2017; Schupp et al. 2021; Tabula Muris et al. 2018) and is also induced by type I interferon (Mar et al. 2018). LY6E belongs to the LY6/uPAR superfamily of secreted or plasma membrane-associated proteins that are characterized by a conserved, three-finger folding motif that is maintained through disulfide bonds (Lee et al. 2013). LY6E modulates immune response signaling pathways (Saitoh et al. 1995; Mao, Hunt, and Cheng 2010; Xu et al. 2014) and may be involved in myeloid and lymphoid cell development (Classon and Coverdale 1994; Wu et al. 1995; Noda et al. 1996). The murine ortholog of LY6E was also identified as the cognate receptor for mouse syncytin-A (Bacquin et al. 2017), which mediates cell to cell fusion during formation of the syncytiotrophoblast layer I of the fetomaternal placenta. Genetic deletion of germline *Ly6e* is incompatible with embryonic development (Zammit et al. 2002) due to disrupted placental architecture (Langford et al. 2018).

The role of LY6E in modulating infection by enveloped RNA viruses has been an active area of investigation over the past decade. LY6E was originally identified through high-throughput screening as an ISG that enhances infection by flaviviruses, a subset of alphaviruses, and influenza A virus (Krishnan et al. 2008; Schoggins et al. 2011; Schoggins et al. 2014). In following studies, LY6E was found to enhance post-attachment viral entry of flaviviruses and influenza A virus (Mar et al. 2018; Hackett and Cherry 2018) as well as HIV-1 (Yu, Liang, and Liu 2017). In more recent screening efforts, LY6E was identified as a restriction factor for coronaviruses (Pfaender et al. 2020; Wickenhagen et al. 2021; Danziger et al. 2022; Mac Kain et al. 2022) including members of the betacoronavirus genus: mouse hepatitis virus (MHV) and severe acute respiratory syndrome coronavirus 2 (SARS-CoV-2). However, our understanding of how LY6E controls coronavirus pathogenesis *in vivo* has only been demonstrated with MHV (Pfaender et al. 2020), due to previous limitations of available mouse models for studying SARS-CoV-2 infection.

In this study, we sought to address two principal knowledge gaps: 1) the contribution of *Ly6e* in distinct immune cell subsets to controlling MHV infection in mice, and 2) the role of *Ly6e* in an immunocompetent murine model of non-mouse-adapted SARS-CoV-2 infection without ectopic expression of human *ACE2*. For the MHV studies, we used comparative models of intraperitoneal versus intranasal administration in i) new mouse lines with *Ly6e* knocked out in distinct immune cells, ii) bone-marrow chimeric mice, and iii) *Ly6e* whole body knockouts. We found that *Ly6e*-dependent control of MHV-induced pathogenesis is cumulatively conferred by *Lyz2*-positive cells, by a radioresistant *Vav1*-positive cell population, and by non-hematopoietic *Vav1*-negative cells. We additionally found that whole body *Ly6e* knockout mice are more susceptible to SARS-CoV-2, with pathogenic outcomes that mirror mild to moderate COVID-19 disease in humans. From transcriptomic analysis of lungs of SARS-CoV-2 infected animals, we found a gene signature consistent with loss of secretory cell-associated transcripts, which may underly the clinical disease observed after *Ly6e* deletion.

## Results

### *Ly6e* in *Vav1*-expressing cells protects against peripheral and systemic coronavirus infection

We previously generated a LY6E-deficient mouse model in which LY6E expression was ablated in all immune cells by crossing a Vav1-iCre transgenic mouse with mice bearing “floxed” *Ly6e* alleles (*Ly6e^fl/fl^*) (Pfaender et al. 2020). Approximately 75% of the mice resulting from this cross, *Ly6e^ΔVav1^*, succumbed to systemic (intraperitoneal) infection with 50,000 PFU MHV, whereas control *Ly6e^fl/fl^* mice survived. To examine how *Ly6e* expression protects mucosal barriers, we used the same mice to compare disease and mortality after MHV infection via peripheral (intranasal) or systemic (intraperitoneal) routes of infection. Mice were infected with 5,000 PFU of MHV (polytropic strain A59) by either route and monitored daily for survival. *Ly6e^fl/fl^* mice were mostly resistant to coronavirus infection by both intranasal (Figure 1A) and intraperitoneal (Figure 1G) routes. *Ly6e^ΔVav1^* mice exhibited 100% lethality when infected intranasally (Figure 1A) but were comparatively less sensitive to intraperitoneal infection, with a 56% mortality rate (Figure 1G).

**Figure 1.**
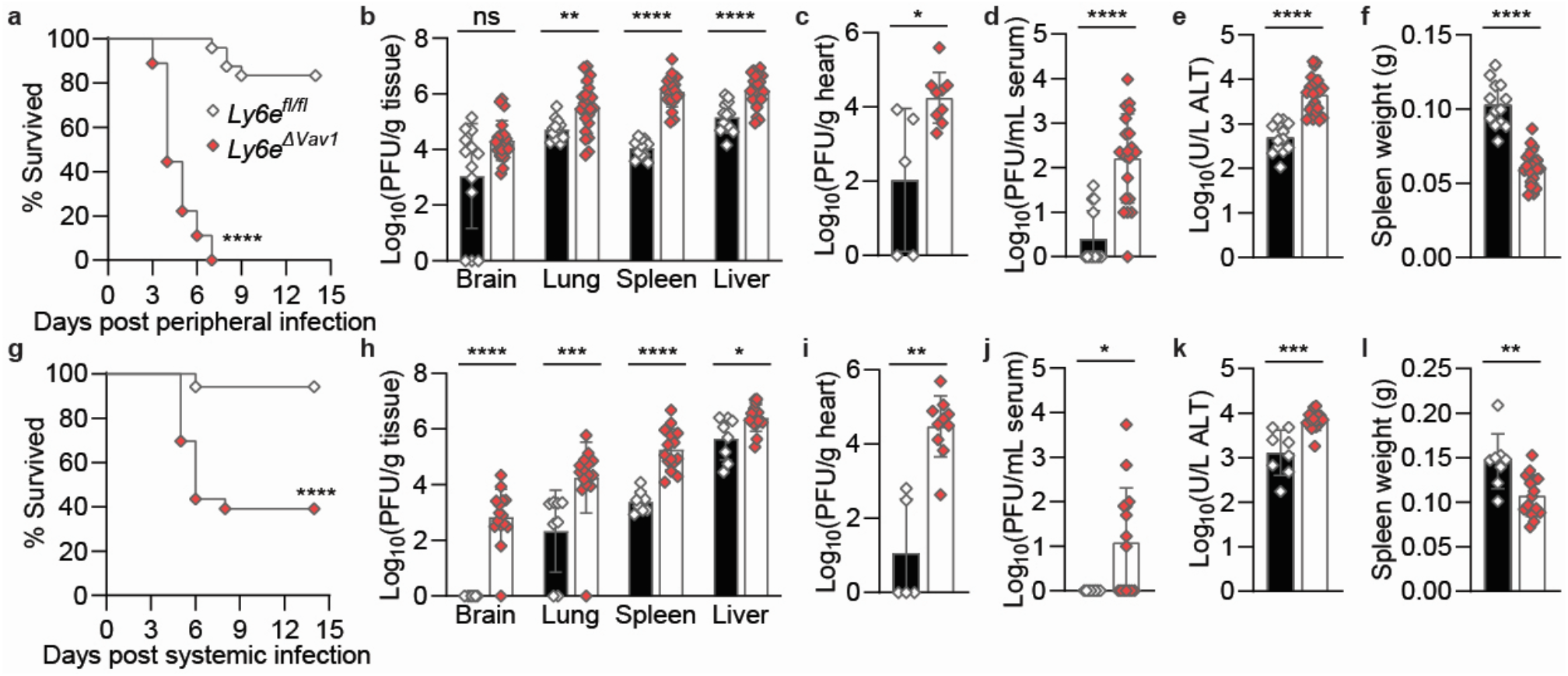
Ly6e in Vav1-expressing cells is critical for viral control in peripheral and systemic coronavirus infections. (A-F) Ly6e^fl/fl^ and Ly6e^ΔVav1^ mice were peripherally (intranasally or IN) infected with 5,000 PFU MHV-A59 and assessed for (A) survival (n=24 Ly6e^fl/fl^ and n=18 Ly6e^ΔVav1^), (B) viral burden in brain, lung, spleen, and liver (n=13 Ly6e^fl/fl^ and n=21 Ly6e^ΔVav1^), (C) viral burden in heart (n=5 Ly6e^fl/fl^ and n=9 Ly6e^ΔVav1^), (D) viral burden in serum, (E) serum alanine aminotransferase, and (F) post-mortem spleen weight (n=13 Ly6e^fl/fl^ and n=21 Ly6e^ΔVav1^ for D-F). (G-L) Ly6e^fl/fl^ and Ly6e^ΔVav1^ were systemically (intraperitoneally or IP) infected with 5,000 PFU MHV-A59 and assessed for (G) survival (n=34 Ly6e^fl/fl^ and n=23 Ly6e^ΔVav1^), (H) viral burden in brain, lung, spleen, and liver (n=8 Ly6e^fl/fl^ and n=15 Ly6e^ΔVav1^), (I) viral burden in heart (n=5 Ly6e^fl/fl^ and n=10 Ly6e^ΔVav1^), (J) viral burden in serum, (K) serum alanine aminotransferase, and (L) post-mortem spleen weight (n=8 Ly6e^fl/fl^ and n=15 Ly6e^ΔVav1^ for J-L). Male and female mice were used at an approximately 1 to 1 ratio for these experiments. Statistical significance was determined by log-rank (Mantel-Cox) tests (A, G), Mann-Whitney test (B-D, H-J), and unpaired t test (E-F, K-L). Error bars represent mean ± standard deviation. ns p>0.05, * p<0.05, ** p<0.01, *** p<0.001, **** p<0.0001.

To evaluate how *Ly6e* in *Vav1*-expressing cells controls viral replication and pathogenesis, we examined viral burden and disease outcomes in peripherally infected mice at 4 days post-infection and in systemically infected mice at 5 days post-infection. We chose a 4-day time point due to earlier onset of lethal disease after intranasal inoculation (Figure 1A) relative to after intraperitoneal inoculation (Figure 1G). Peripherally infected *Ly6e^ΔVav1^* mice had increased viral burden in the lung compared to *Ly6e^fl/fl^* littermates, demonstrating that *Ly6e* in *Vav1*-expressing cells contributes to control of mucosal coronavirus infection of the airways (Figure 1B). Murine coronavirus infection of the brain, which occurs after intranasal infection via the olfactory bulb (Cupovic et al. 2016), was not statistically different between *Ly6e^fl/fl^* and *Ly6e^ΔVav1^* mice (Figure 1B). *Ly6e^ΔVav1^* mice had elevated viral titers in spleen, liver (Figure 1B), heart (Figure 1C), and serum (Figure 1D), indicating that *Ly6e* limited viral dissemination beyond the respiratory tract or replication within those organs. Serum alanine aminotransferase (ALT) levels, which indicates hepatic damage, were higher in *Ly6e^ΔVav1^* mice when compared to *Ly6e^fl/fl^* littermates (Figure 1E). Average post-mortem spleen size was significantly lower in *Ly6e^ΔVav1^* mice (Figure 1F).

Systemic coronavirus infection via intraperitoneal injection resulted in higher viral titers in visceral organs, including lung, spleen, liver (Figure 1H), and heart (Figure 1I), of *Ly6e^ΔVav1^* mice when compared to *Ly6e^fl/fl^* littermates. Strikingly, infectious virus was recovered from the brains of 14 out of 15 intraperitoneally infected *Ly6e^ΔVav1^* mice but was undetectable in the brains of *Ly6e^fl/fl^* littermates (Figure 1H). Nearly half of the *Ly6e^ΔVav1^* mice also had viremia after intraperitoneal infection (Figure 1J). As observed in our previous study with 50,000 PFU MHV (Pfaender et al. 2020), *Ly6e^ΔVav1^* mice that were systemically infected with 5,000 PFU MHV had elevated serum ALT (Figure 1K). Spleens of *Ly6e^ΔVav1^* mice were moderately smaller than those of their *Ly6e^fl/fl^* littermates (Figure 1L).

Our data demonstrate that *Ly6e* in *Vav1*-expressing cells protects against systemic and peripheral coronavirus challenge by limiting viral replication in visceral organs and dissemination from the initial site of inoculation. In the absence of *Ly6e*, uncontrolled coronavirus infection causes hepatitis and splenic hypoplasia, which may contribute to increased lethality.

### *Ly6e* in *Lyz2*-expressing cells and B cells exert organ-specific control of murine coronavirus infection

As Vav1-iCre affects gene expression in diverse immune cell subsets, we next sought to determine the contribution of distinct immune cell types to the pathogenic phenotypes observed in infected *Ly6e^ΔVav1^* mice. Accordingly, we generated new conditional knockout mouse strains using distinct Cre-driver mice. Ablation of *Ly6e* gene expression in lung alveolar macrophages was confirmed for *Ly6e^ΔLyz2^* and *Ly6e^ΔCD11c^* mice relative to cells isolated from *Ly6e^fl/fl^* mice (Figure 2A)). Targeted *Ly6e* deletion in splenic CD4+ T cells, CD8α+ T cells, and CD19+ B cells was also confirmed for *Ly6e^ΔCD4^, Ly6e^ΔCD8a^*, and *Ly6e^ΔCD19^* mice, respectively (Figure 2A, S1A). Conditional knockout mice were challenged intranasally with 5,000 PFU MHV and monitored daily for survival (Figure 2B). *Ly6e^ΔLyz2^* mice, which we predicted to have targeted loss of *Ly6e* in subsets of macrophages, monocytes, neutrophils, and epithelial cells (Abram et al. 2014; McCubbrey et al. 2017), partially recapitulated the susceptibility of *Ly6e^ΔVav1^* mice to peripheral coronavirus infection. *Ly6e^ΔCD8a^* mice, which are predicted to have loss of *Ly6e* in CD8α+ T cells and CD8α+ dendritic cells, exhibited a modest susceptibility to intranasal coronavirus infection. *Ly6e^ΔCD11c^, Ly6e^ΔCD4^*, and *Ly6e^ΔCD19^* conditional knockout mice were resistant to coronavirus-induced disease after intranasal infection when compared to *Ly6e^fl/fl^* littermates. In contrast to *Ly6e^ΔVav1^* mice which were sensitive to systemic coronavirus infection (Figure 1G), *Ly6e^ΔLyz2^, Ly6e^ΔCD11c^, Ly6e^ΔCD4^, Ly6e^ΔCD8a^*, and *Ly6e^ΔCD19^* were fully resistant to intraperitoneal challenge with 5,000 PFU MHV (Figure S1B).

**Figure 2.**
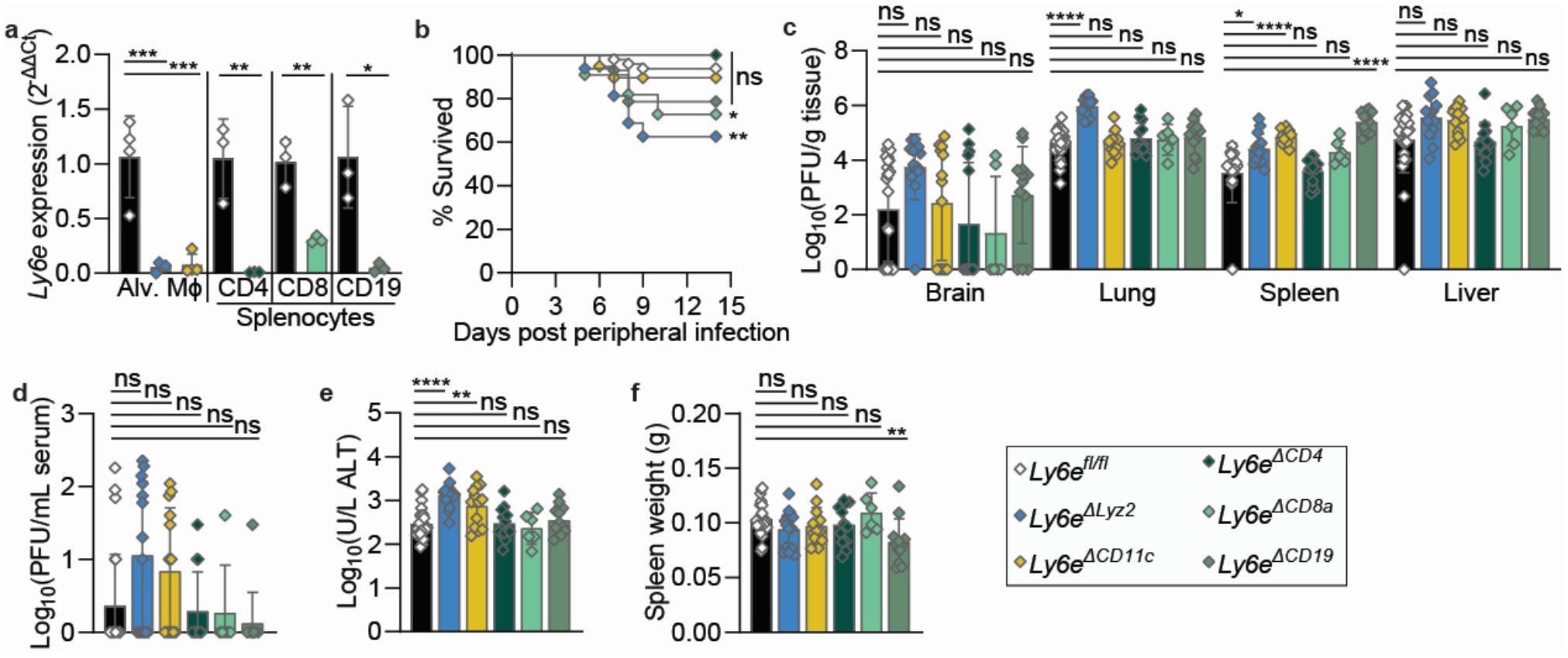
Cell type-specific contributions of Ly6e during peripheral coronavirus infections. (A) Relative Ly6e mRNA levels in lung alveolar macrophages (n=4 Ly6e^fl/fl^, n=4 Ly6e^ΔLyz2^, n=4 Ly6e^ΔCD11c^), splenic CD3ε+CD4+ T cells (n=3 Ly6e^fl/fl^ and n=3 Ly6e^ΔCD4^), splenic CD3ε+CD8α+ T cells (n=3 Ly6e^fl/fl^ and n=3 Ly6e^ΔCD8a^), and splenic CD19+CD3ε-B cells (n=3 Ly6e^fl/fl^ and n=3 Ly6e^ΔCD19^). (B-F) Mice were peripherally (intranasally or IN) infected with 5,000 PFU MHV-A59 and assessed for (B) survival (n=48 Ly6e^fl/fl^, n=16 Ly6e^ΔLyz2^, n=19 Ly6e^ΔCD11c^, n=13 Ly6e^ΔCD4^, n=11 Ly6e^ΔCD8a^, and n=14 Ly6e^ΔCD19^), (C) viral burden in brain, lung, spleen, and liver, (D) viral burden in serum, (E) serum alanine aminotransferase, and (F) post-mortem spleen weight (n=26 Ly6e^fl/fl^, n=14 Ly6e^ΔLyz2^, n=13 Ly6e^ΔCD11c^, n=13 Ly6e^ΔCD4^, n=6 Ly6e^ΔCD8a^, and n=12 Ly6e^ΔCD19^ for C-F). Male and female mice were used at an approximately 1 to 1 ratio for these experiments. Statistical significance was determined by log-rank (Mantel-Cox) tests (B), Kruskal-Wallis test (C-D), and one-way ANOVA (E-F). Error bars represent mean ± standard deviation. ns p>0.05, *p<0.05, ** p<0.01, ***p<0.001, ****p<0.0001.

To determine how loss of *Ly6e* in distinct cell types affects coronavirus replication and dissemination, we determined viral burden in tissues and serum harvested from peripherally infected mice at 4 days post-infection (Figure 2C-D) and systemically infected mice at 5 days post-infection (Figure S1C-D). In intranasally infected mice, viral titers in the brain, liver (Figure 2C), and serum (Figure 2D) were unaffected by *Ly6e* deletion in any compartment. Only peripherally infected *Ly6e^ΔLyz2^* mice had higher viral burden in the lung relative to *Ly6e^fl/fl^* littermates, while *Ly6e^ΔLyz2^, Ly6e^ΔCD11c^, Ly6e^ΔCD19^* mice had increased viral burden of the spleen (Figure 2C). Serum ALT levels were elevated in only *Ly6e^ΔLyz2^* and *Ly6e^ΔCD11c^* mice (Figure 2E). *Ly6e^ΔCD19^* mice also exhibited mild splenic hypoplasia after intranasal infection that was not observed in other conditional knockout strains (Figure 2F).

In systemically infected mice, infectious virus was not detected in brains of any conditional knockout strain (Figure S1C). In contrast to the peripheral infection route, conditional deletion of *Ly6e* in *Lyz2-*expressing cells or any other cell type had no effect on lung viral burden after intraperitoneal infection. Only *Ly6e^ΔCD19^* mice had elevated viral titers in spleen and liver, consistent with our previous finding that Ly6e-deficient splenic B cells are highly permissive to MHV. A subpopulation of systemically infected mice had viremia after intraperitoneal infection, with a modest elevation observed in *Ly6e^ΔCD8a^* mice (Figure S1D). Serum ALT levels were only higher in *Ly6e^ΔLysz2^* and *Ly6e^ΔCD19^* mice after systemic infection (Figure S1E). In contrast to *Ly6e^ΔVav1^* mice, which exhibited splenic hypoplasia after systemic coronavirus infection (Figure 1L), *Ly6e^ΔLyz2^, Ly6e^ΔCD11c^*, and *Ly6e^ΔCD8a^* mice had larger spleens relative to *Ly6e^fl/fl^* littermates (Figure S1F).

### *Ly6e* in a radioresistant *Vav1*-expressing cellular compartment limits coronavirus-mediated disease

None of the conditional knockout strains described above fully recapitulated the pathogenesis phenotypes of *Ly6e^ΔVav1^* mice. Accordingly, we hypothesized that a *Ly6e*-deficient non-immune cell compartment generated by the Vav1-iCre transgene may contribute to the phenotype (Joseph et al. 2013; Siegemund et al. 2015). To test this model, we generated bone marrow chimeric mice (*Ly6e^fl→ΔVav1^*) to reconstitute *Ly6e* expression in the hematopoietic compartment of *Ly6e^ΔVav1^* mice (Figure 3A). We also generated control mice (*Ly6e^fl→fl^*) by transplanting *Ly6e*-sufficient bone marrow into irradiated *Ly6e^fl/fl^* recipients. Restoration of *Ly6e* expression in the spleen (Figure 3B) and in CD45+ immune cells from the lung (Figure 3C, S2A) was confirmed in *Ly6e^fl→ΔVav1^* chimeras. Vav1-iCre mice have been reported to express iCre in endothelial cells (Joseph et al. 2013); however, we found that CD31+CD45-EpCam-endothelial cells isolated from the lungs of *Ly6e^ΔVav1^* mice had similar *Ly6e* expression as *Ly6e^fl/fl^* littermates (Figure 3D, S2A). Finally, we analyzed the relative composition of donor (CD45.1+) and recipient (CD45.2+) immune cells in lung, spleen, and blood in the *Ly6e^fl→fl^* and *Ly6e^fl→ΔVav1^* chimeras and found that donor cells respectively comprised an average of 95.4%, 97.1%, and 96.7% of the total CD45+ cellular compartment (Figure S2B-C).

**Figure 3.**
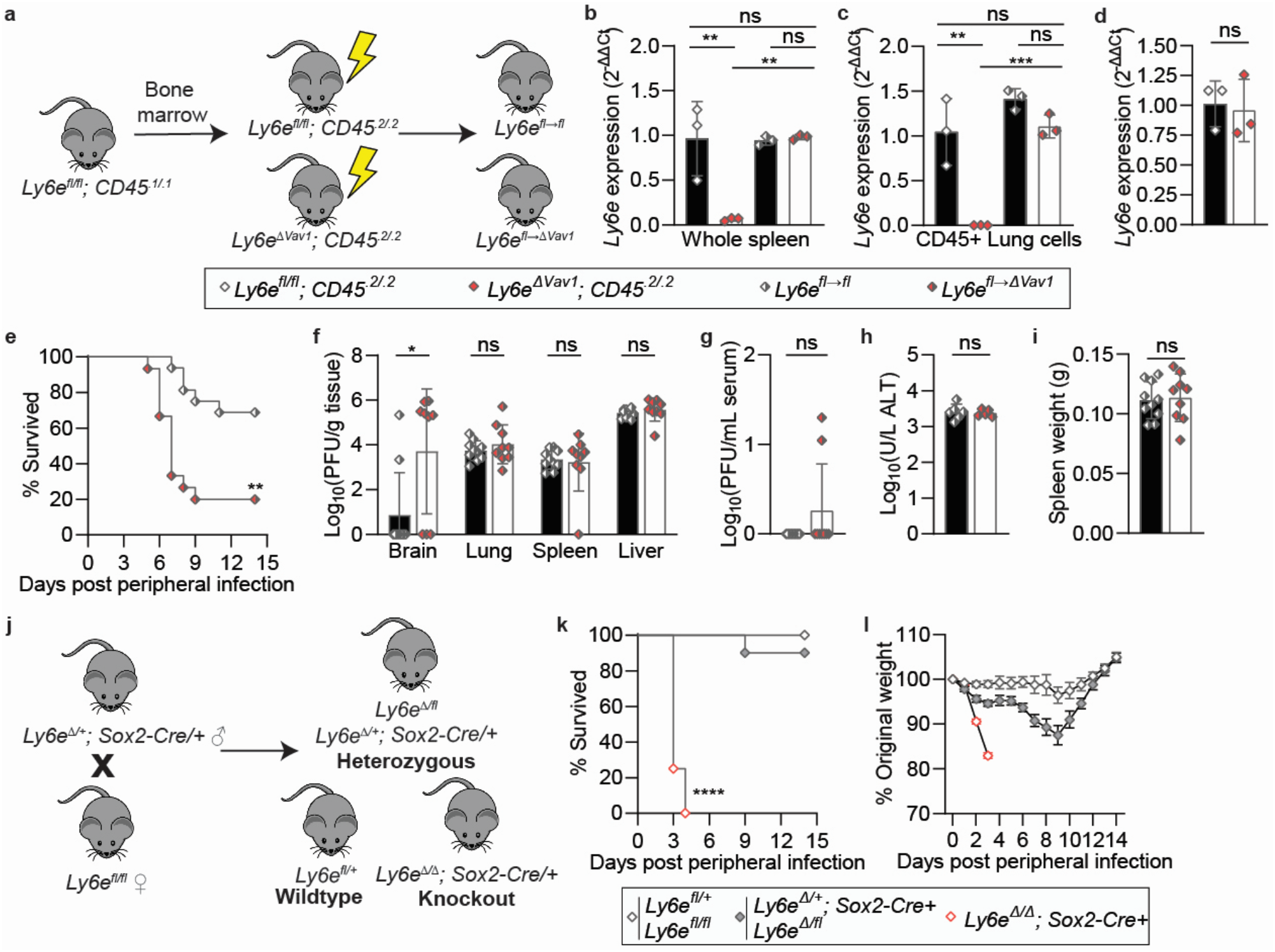
Reconstitution of Ly6e expression in the hematopoietic compartment reveals that Ly6e in radioresistant Vav1-expressing cells is protective against peripheral and systemic coronavirus infection. (A) Experimental scheme to generate bone marrow chimeric mice. (B-D) Relative Ly6e mRNA levels in (B) whole spleen, (C) lung CD45+ immune cells (n=3 Ly6e^fl/fl^, n=3 Ly6e^ΔVav1^, n=3 Ly6e^fl→fl^, n=3 Ly6e^fl→ΔVav1^ for B-C), and (D) lung CD31+CD45-EpCam-endothelial cells (n=3 Ly6e^fl/fl^ and n=3 Ly6e^ΔVav1^). (E-I) Ly6e^f→fl^ and Ly6e^fl→ΔVav1^ mice were peripherally (intranasally or IN) infected with 1,000 PFU MHV-A59 and assessed for (E) survival (n=16 Ly6e^f→fl^ and n=15 Ly6e^fl→ΔVav1^), (F) viral burden in brain, lung, spleen, and liver, (G) viral burden in serum, (H) serum alanine aminotransferase, and (I) post-mortem spleen weight (n=10 Ly6e^f→fl^ and n=9 Ly6e^fl→ΔVav1^ for F-I). (J) Breeding scheme to generate whole body Ly6e heterozygous and knockout mice. (K-L) Ly6e wildtype, heterozygous, and knockout mice were peripherally (IN) infected with 5,000 PFU MHV-A59 and assessed for (K) survival and (L) daily weight loss (n=6 wildtype, n=10 heterozygous, n=10 knockout). Male and female mice were used at an approximately 1 to 1 ratio for these experiments. Statistical significance was determined by log-rank (Mantel-Cox) tests (E, K), Mann-Whitney test (F-G), one-way ANOVA (B-C), and unpaired t test (D, H-I). Error bars represent mean ± standard deviation in all panels except for (L) where error bars indicate mean ± SEM. ns p>0.05, *p<0.05, **p<0.01, ***p<0.001, ****p<0.0001.

*Ly6e^fl→fl^* and *Ly6e^fl→ΔVav1^* chimeras were intranasally challenged with 1,000 PFU MHV and monitored daily for survival. We reduced the dose of viral inoculum due to the increased sensitivity of the chimeras to 5,000 PFU MHV, regardless of recipient genotype (data not shown). While onset of lethal disease was delayed relative to non-chimeric *Ly6e^ΔVav1^* mice (Figure 1A), *Ly6e^fl→ΔVav1^* mice succumbed at a greater rate to mucosal infection than their *Ly6e^fl→fl^* littermates (Figure 3E). Viral burden was assessed in chimeric mice 6 days after peripheral infection, to account for the delay in disease onset relative to intranasally infected *Ly6e^ΔVav1^* mice. In contrast to *Ly6e^ΔVav1^* mice (Figure 1B), *Ly6e^fl→ΔVav1^* mice exhibited similar viral burden in lung, spleen, and liver as *Ly6e^fl→fl^* littermates (Figure 3F). Strikingly, high viral load was detected in the brains of 6 of the 9 *Ly6e^fl→ΔVav1^* mice, but only in 2 of the 10 *Ly6e^f→fl^* control mice. Viremia was observed in several *Ly6e^fl→ΔVav1^* mice at this time point but was not significantly elevated compared to control mice (Figure 3G). Serum ALT levels (Figure 3H) and spleen weight (Figure 3I) were the same in *Ly6e^fl→ΔVav1^* mice as *Ly6e^fl→fl^* mice, indicating that *Ly6e*-sufficient hematopoietic donor cells rescued hepatic damage and splenic hypoplasia.

Together, our bone marrow chimera studies show that *Ly6e* expressed by a radioresistant *Vav1-*expressing cell type is important for resistance to peripheral coronavirus infection. However, the contribution to viral control of other *Ly6e*-expressing cell that do not express *Vav1* is unclear. To address this gap in knowledge, we used Sox2-Cre transgenic mice to generate *Ly6e* whole body knockout mice (Figure 3J). This strategy bypasses the embryonic lethal effects of germline *Ly6e* deletion (Zammit et al. 2002) by preserving gene expression in the placenta (Langford et al. 2018). *Ly6e* whole body knockout mice (*Ly6e^Δ/Δ^; Sox2-Cre*+) succumbed rapidly to intranasal challenge with 5,000 PFU MHV (Figure 3K). *Ly6e* heterozygous mice exhibited transient weight loss (Figure 3L) but ultimately survived peripheral coronavirus infection at a similar rate as wildtype animals. The additional loss of *Ly6e* in *Vav1*-negative cell types accelerated coronavirus-driven lethality compared to genetic ablation in *Vav1*-expressing cell types alone (Figure 1A), demonstrating a major contribution of the non-hematopoietic compartment to surviving murine coronavirus challenge.

### *Ly6e* restrains pathogenesis of a SARS-CoV-2 variant of concern in mice

Ancestral strains of SARS-CoV-2 could not infect mice without transgenic overexpression of human ACE2 (Bao et al. 2020) or reverse genetic manipulation and passaging in mice (Dinnon et al. 2020; Leist et al. 2020). This hindered our ability to examine infection of clinical isolates of SARS-CoV-2 in LY6E-deficient mice. At the end of 2020 through early 2021, several SARS-CoV-2 variants of concern with the N501Y mutation in the receptor binding domain of the spike protein, which has been associated with increased affinity for murine ACE2 (Gu et al. 2020), began to circulate globally. The SARS-CoV-2 Gamma variant (P.1), which was originally detected in January 2021, was shown to replicate efficiently in the respiratory tracts of young C57BL/6J mice without causing clinically observable disease such as weight loss (Imai et al. 2021). Viral antigen was detected in the respiratory bronchial and alveolar epithelial cells of infected mice, indicating the P.1 replicates in non-hematopoietic cells in the airways. The molecular and cellular pathways that confer murine resistance to SARS-CoV-2 variants such as P.1 remain undefined.

We hypothesized that *Ly6e*, which is constitutively expressed at high levels in most tissues (Tabula Muris et al. 2018) and lung cell types (Schupp et al. 2021), contributes to murine resistance to SARS-CoV-2 infection. Sox2-driven Cre recombinase efficiently deleted *Ly6e* expression in multiple tissues from *Ly6e* homozygous knockout mice, including the lungs (Figure S3A). *Ly6e* wildtype, heterozygous, and knockout mice were challenged intranasally with 60,000 PFU P.1 SARS-CoV-2 and monitored daily for weight loss and survival. On day 2 post-infection, knockout mice uniformly lost 10% of their original body weight while wildtype and heterozygous mice displayed little to no weight loss (Figure 4A). All knockout mice began regaining weight by day 4, and by day 9 post-infection all mice had fully recovered. Decreasing the viral inoculum by more than 6-fold to 8,700 PFU did not alter the disease course of *Ly6e* wildtype or knockout mice (Figure 4B). At 3 days post-infection, lung viral burden was elevated in *Ly6e* knockout mice compared to wildtype mice, as measured by plaque assay (Figure 4C) and quantitative PCR (Figure 4D). To further understand the role of antiviral effectors in murine resistance to SARS-CoV-2, we intranasally challenged mice lacking type I interferon (*Ifnar*^-/-^) and type III interferon (*Ifnlr*^-/-^) signaling with 60,000 PFU P.1 SARS-CoV-2 and compared weight change to infected wildtype C57BL/6J mice. Strikingly, *Ifnar*^-/-^ and *Ifnlr*^-/-^ mice displayed similar resistance as wildtype mice to SARS-CoV-2 induced weight loss (Figure 4E). Despite no clinical signs of disease, the absence of type I interferon and type III interferon signaling in these mice increased viral replication in the airways at 3 days post-infection (Figure 4F-G). Collectively, our data suggests that interferons contribute to limiting SARS-CoV-2 replication in the lungs, while *Ly6e* helps to control early viral pathogenesis before other components of the antiviral immune response ultimately clear infection and accelerate host recovery.

**Figure 4.**
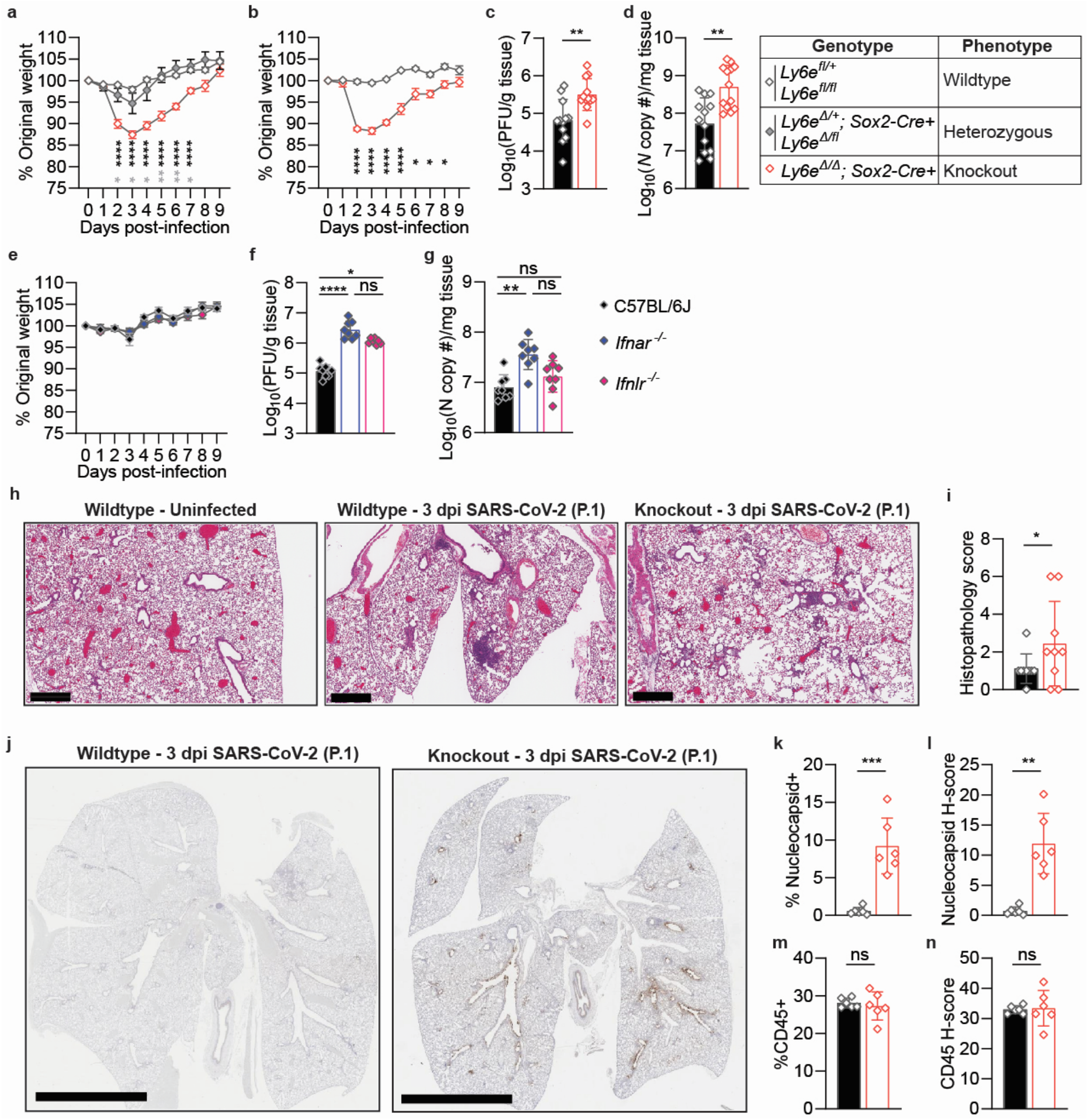
Ly6e reduces SARS-CoV-2 replication and pathogenesis in mice. Ly6e wildtype, heterozygous, and knockout mice were intranasally (IN) infected with (A) 60,000 PFU P. 1 SARS-CoV-2 (n=7 wildtype, n=4 heterozygous, and n=7 knockout) or (B) 8,700 PFU P. 1 SARS-CoV-2 (n=5 wildtype and n=5 knockout) and monitored for daily weight loss. Ly6e wildtype and knockout mice were IN infected with 60,000 PFU P. 1 SARS-CoV-2 and viral burden in lung was measured by (C) plaque assay (n=12 wildtype and n=12 knockout) and (D) quantitative PCR (n=13 wildtype and n=13 knockout) 3 days post-infection. C57BL/6J, Ifnar^-/-^, and Ifnar^-/-^ mice were IN infected with 60,000 PFU P. 1 SARS-CoV-2 and monitored for (E) daily weight loss (n=10 C57BL6/J, n=10 IFNAR-/-, and n=10 IFNLR^-/-^) and lung viral burden as measured by (F) plaque assay and (G) quantitative PCR (n=8 C57BL6/J, n=8 Ifnar^-/-^, and n=8 Ifnlr^-/-^ for F-G) 3 days post-infection. (H) Lung sections stained with hematoxylin and eosin. (I) Histopathological scoring for features of acute pneumonia in lungs from SARS-CoV-2 infected wildtype (n=9) and knockout (n=9) mice at 3 days post-infection. (J) Representative whole lung sections stained for SARS-CoV-2 nucleocapsid (DAB) and a hematoxylin counterstain. Analyses for K-N was performed on lung sections from SARS-CoV-2 infected wildtype (n=6) and knockout (n=6) mice. Percentage of SARS-CoV-2 nucleocapsid-positive cells of total lung cells (K) and nucleocapsid stain H-score which integrates staining intensity and proportion of total cells (L). Percentage of CD45-positive cells of total lung cells (M) and CD45 stain H-score which integrates staining intensity and proportion of total cells (N). For (A-G), male and female mice were used at an approximately 1 to 1 ratio. For (H-N), males and female mice were used at an 8:1 ratio based on availability. Statistical significance was determined by multiple unpaired t tests (A-B, K, M), Mann-Whitney test (C-D, F-G, L, N), and Kolmogorov-Smirnov test (I). In (A-B), black symbols indicate statistical comparison of wildtype and knockout mice, and gray symbols indicate comparison of heterozygous and knockout mice (A). Error bars represent mean ± standard deviation in all panels except for A, B, and E where error bars indicate mean ± SEM. ns p>0.05, * p<0.05, ** p<0.01, ***p<0.001, **** p<0.0001. Scale bars: 600μm (H) and 5mm (J).

To gain further insight into how *Ly6e* controls early viral pathogenesis, we performed histological analyses of lung 3 days post-infection. SARS-CoV-2 infection caused mild perivascular inflammation in most wildtype mice, whereas knockout mice exhibited moderate perivascular inflammation in most animals and mild pleural and alveolar inflammation in several animals. Four of the nine *Ly6e* knockout mice also exhibited signs of hemorrhage that was not observed in wildtype animals (Figure 4H-I, Table S1). Lung sections from infected animals were also probed with an antibody that detects SARS-CoV-2 nucleocapsid protein (Figure 4J). Immunohistochemistry analysis indicated immunopositivity in the epithelium of the main airways of all *Ly6e* knockout mice, particularly in cells sloughed off into the airway, with occasional immunopositive staining in alveolar epithelial cells and intra-alveolar macrophages. Unbiased quantification of nucleocapsid-containing cells revealed that significantly more cells in the lungs of *Ly6e* knockout mice were productively infected with virus (Figures 4K-L, S3B, Table S2). Infiltration of CD45+ immune cells in response to SARS-CoV-2 infection was similar between infected wildtype and *Ly6e* knockout mice (Figure 4M-N, S3C, Table S2). Our data suggests that *Ly6e* expression helps restrain SARS-CoV-2-induced lung pathology that appears independent of inflammatory immune cell recruitment.

### Loss of Ly6e correlates with alterations in pulmonary homeostatic gene expression

We next sought to uncover the cause of clinical signs of disease in SARS-CoV-2-infected *Ly6e* knockout mice through transcriptomic analysis of lung tissue. In the absence of infection, transcriptomic analyses of whole lungs from *Ly6e* wildtype and *Ly6e* knockout mice revealed only 30 genes whose mRNA levels were significantly changed (*p_adj_* < 0.05) more than two-fold in the absence of *Ly6e* expression (Figure 5A, 5D). By contrast, a robust induction of numerous interferon-stimulated genes and mucosal barrier-associated genes (*Reg3g, Saa3*) was observed in lungs from SARS-CoV-2 infected wildtype mice when compared to uninfected wildtype animals (Figure 5B, 5E). *Ly6e* knockout mice exhibited a similar interferon-stimulated gene signature after SARS-CoV-2 infection when compared to uninfected knockout animals (Figure 5C, 5F). We next compared the expression levels of the top 25 genes induced by SARS-CoV-2 infection in wildtype mice relative to *Ly6e* knockout mice. Interferon-stimulated genes, for the most part, were not differentially induced by SARS-CoV-2 infection in wildtype or knockout mice (Figure 5G). The gene *Muc5b*, which encodes a respiratory tract mucin glycoprotein that contributes to mucociliary clearance and antibacterial defense (Roy et al. 2014) was significantly elevated by an average of two-fold in the lungs of SARS-CoV-2 infected wildtype mice relative to infected knockout mice. A polymorphism that increases expression of human *MUC5B* in the lungs was identified by three separate meta-analysis studies as protective against severe COVID-19 (Fadista et al. 2021; van Moorsel et al. 2021; Verma et al. 2021), indicating that induction of *Muc5b* by SARS-CoV-2 may help control respiratory disease. Other secretory cell-associated genes, such as *Muc5ac, Scgb1a1*, and *Scgb3a1* (Schupp et al. 2021), were elevated in the lungs of infected, wildtype mice relative to infected, knockout mice (Figure 5H). A subset of secretory cell genes (*Muc5b, Pigr, Bpifb1, Retnla, Muc5ac*) that were elevated in infected wildtype mice relative to infected knockout mice were significantly induced 3.5 to twelve-fold by SARS-CoV-2 infection in wildtype animals (Figure S4A). In contrast, the same subset of genes was induced only up to 2.3-fold by SARS-CoV-2 infection in knockout mice, and a separate subset of genes that are expressed highly in secretory cells and to a lesser degree in ciliated cells (*Scgb1a1, Cyp2f2, Cldn10*) were decreased by 4 to 17-fold after SARS-CoV-2 infection (Figure S4B). Intriguingly, gene expression of the mouse ortholog of the SARS-CoV-2 receptor ACE2 is highly enriched in respiratory tract secretory cells relative to other lung cell subtypes whereas *Ly6e* is ubiquitously detected irrespective of cell type (Schupp et al. 2021). Gene Ontology (GO) enrichment analysis of genes differentially expressed between SARS-CoV-2 infected wildtype and knockout mice revealed signatures of increased leukocyte motility and adhesion in wildtype animals (Figure 5I). Collectively, our data suggests that *Ly6e* expression may protect lung secretory cells from SARS-CoV-2 infection, which in turn preserves protective *Muc5b* expression and pulmonary leukocyte mobility.

**Figure 5.**
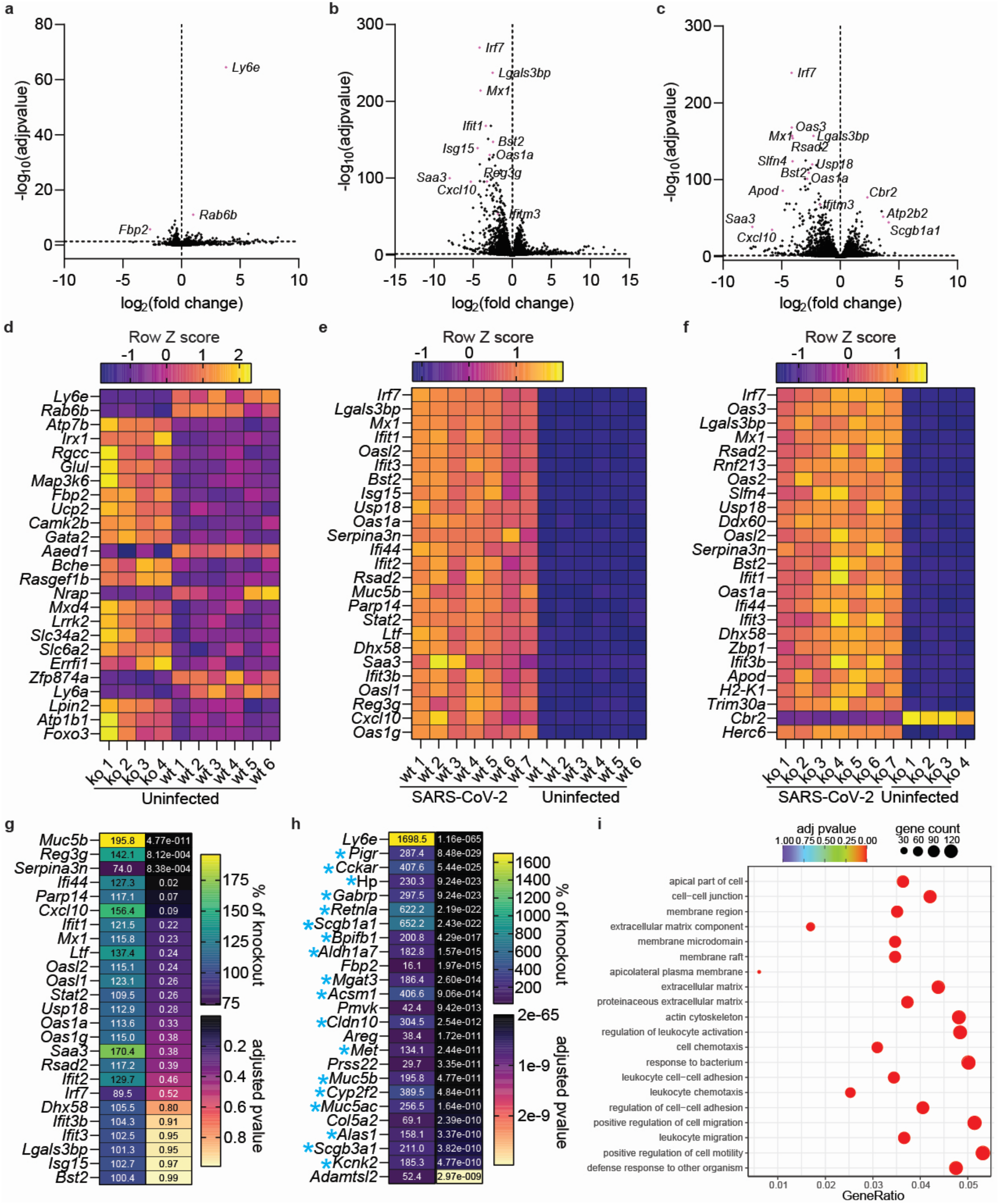
Ly6e expression preserves a secretory cell gene signature after SARS-CoV-2 infection. Ly6e wildtype and knockout mice were intranasally (IN) infected with 60,000 PFU P. 1 SARS-CoV-2 and euthanized 3 days post-infection for whole lung RNA isolation and subsequent mRNA sequencing. (A-C) Volcano plot summarization of differentially expressed genes (DEG) between uninfected Ly6e wildtype (n=6) and Ly6e knockout (n=4) mice (A), uninfected wildtype (n=6) and SARS-CoV-2 infected wildtype (n=7) mice (B), and uninfected knockout (n=4) and SARS-CoV-2 infected knockout (n=7) mice (C). (D-F) Z-score heatmaps of the top 25 most significantly differentially expressed genes respectively for data summarized in A-C. (G-H) Relative difference between SARS-CoV-2 infected wildtype and Ly6e knockout mice in the top 25 genes induced by viral infection in wildtype mice (G) and the top 25 most significantly differentially expressed genes (H). Genes expressed highly by secretory cells are indicated by a blue asterisk. (I) Gene Ontology (GO) enrichment analysis of transcriptomic data from SARS-CoV-2 infected wildtype and knockout mice. The abscissa is the ratio of the number of differential genes linked with the GO term to the total number of differential genes. The size of a point represents the number of genes annotated to a specific GO term, and the color from red to purple represents the significant level of the enrichment. Male and female mice were used at an approximately 1 to 1 ratio for these experiments. Determination of statistical significance is described in the Methods section titled ‘RNA-Seq.’

## Discussion

The cell type-specific contribution of individual cell-intrinsic antiviral effectors to controlling coronavirus pathogenesis has not been previously investigated. Our study expands our understanding of *Ly6e*-mediated control of murine coronavirus infection by identifying specific cellular compartments that require *Ly6e* to establish host resistance. By using a SARS-CoV-2 variant that infects mice without ectopic overexpression of human *ACE2*, we also identified a putative protective role for *Ly6e* in secretory cells which may restrain respiratory disease through production of MUC5B and other factors that control leukocytes.

Our conditional knockout data demonstrate that host control of MHV infection is due in part to unique, organ-specific contributions of *Ly6e* in distinct cellular compartments (Figure 2, S1). *Ly6e* in *Lyz2*-expressing cells limited viral replication in lung and spleen only in the peripheral infection model, suggesting a role for LY6E in constraining respiratory tract replication and dissemination beyond the airways that may contribute to lethal disease. In both peripheral and systemic infection models, serum ALT in *Ly6e^ΔLyz2^* mice was higher, indicating that *Ly6e* may protect a *Lyz2*-expressing cell type that is crucial for limiting coronavirus-driven hepatic damage. *Ly6e* in *CD11c*-expressing cells was dispensable for surviving peripheral and systemic coronavirus challenge but helped control hepatic pathology after intranasal infection only. Elevated splenic viral burden in *Ly6e^ΔCD11c^* mice was observed after intranasal infection but not after intraperitoneal infection, which suggests that *Ly6e* in CD11c+ cells may help control viral dissemination from the airways to the spleen. *Ly6e* in CD19+ B cells, which we previously showed as important for protecting B cells from direct MHV infection and for maintaining splenic and hepatic B cell numbers after intraperitoneal infection (Pfaender et al. 2020), was dispensable for surviving peripheral and systemic coronavirus challenge. In corroboration with our previous study, we showed that *Ly6e* in CD19+ B cells helps control coronavirus replication in spleen in both peripheral and systemic infection models as well as in liver during systemic infection only. LY6E in CD4+ T cells and CD8α+ T cells was dispensable for viral control in most assays, with *Ly6e^ΔCD8a^* mice exhibiting a modest increase in sensitivity to intranasal coronavirus infection. LY6E expression is not essential for protecting T cells from direct MHV infection (Pfaender et al. 2020), but may be involved in regulation of T cell receptor signaling (Saitoh et al. 1995). Cumulatively, our approach to identify which distinct cellular compartments in the total population of *Vav1-*expressing cells require LY6E to control coronavirus-mediated disease revealed a role in *Lyz2*-expressing cells but failed to uncover a single cell type co-expressing *Ly6e* and *Vav1* that could fully recapitulate the phenotype observed in *Ly6e^ΔVav1^* mice (Figure 1A, G).

Our bone marrow chimera studies suggest that LY6E in a radioresistant *Vav1*-expressing cell type contributes significantly to host defense against coronaviruses (Figure 3). We confirmed that *Ly6e* expression was restored in lung and spleen CD45+ cells and that donor CD45.1+ cells almost fully occupy the hematopoietic niche in 3 different compartments. However, several cell types may still retain recipient genetics due to resistance to irradiation, such as microglia in the brain, which highly express *Ly6e* (Tabula Muris et al. 2018). We showed that the pulmonary endothelial cells as a bulk population are unaffected by Vav1-iCre-driven knockout of *Ly6e* (Figure 3D), but it remains unclear whether other endothelial cell subtypes in different organs may have *Vav1*-driven expression of Cre recombinase as suggested by others (Joseph et al. 2013). The MHV receptor CEACAM1 is highly expressed on the endoluminal pole of endothelial cells of cerebral blood vessels that are resistant to coronavirus infection, which consequently prevents viral spread into the brain after intraperitoneal infection (Godfraind et al. 1997). In contrast, we observed viral dissemination into the brain after intraperitoneal infection in *Ly6e^ΔVav1^* animals, but not any other conditional knockout strain, indicating that some *Vav1*-expressing cell type or combination of cell types that was not interrogated in our other approaches may contribute to host resistance to nervous system infection and survival after systemic challenge (Figure 1). This same cell type may also help control viral replication in the brains of intranasally infected bone marrow chimeras (Figure 3). We speculate that microglia, which are not uniformly targeted by the LysM-Cre transgene (Orthgiess et al. 2016) but highly express *Vav1*, are more likely to be a relevant cell type in *Ly6e^ΔVav1^* and *Ly6e^fl→ΔVav1^* mice than brain endothelial cells, which are poor expressors of *Vav1* (Tabula Muris et al. 2018).

Our study also highlights different antiviral mechanisms used to resist a coronavirus that naturally infects mice (MHV) versus a human coronavirus that recently gained tropism for *Mus musculus* (SARS-CoV-2). We showed that whole body expression of *Ly6e* was indispensable for surviving murine coronavirus infection (Figure 3P). Survival of intranasal murine coronavirus infection is also dependent on intact type I and III interferon signaling (Sharma et al. 2022). In contrast, *Ly6e* and type I and III interferon signaling were each dispensable for surviving intranasal challenge with SARS-CoV-2. It is possible that each of these antiviral systems compensate for each other if one is lost. This can be tested in future studies combining Ly6e-defiency with complete loss of interferon signaling, by STAT1 ablation for example.

Ly6e has a clear role in protecting mice from SARS-CoV-2-induced pathogenesis (Figure 4). Ly6e-deficient mice exhibit a non-lethal clinical disease that may potentially model “mild to moderate” COVID-19 in humans, defined by clinical presence of lower respiratory tract disease (Gandhi, Lynch, and Del Rio 2020). Thus, this model may have unique uses for testing vaccines and antivirals, as it recapitulates a more common form of human disease than the more severe and lethal models of mouse-adapted SARS-CoV-2 (Dinnon et al. 2020; Leist et al. 2020; Gu et al. 2020; Huang et al. 2021; Sun et al. 2021; Muruato et al. 2021) or SARS-CoV-2 infection in K18-hACE2 mice (Winkler et al. 2020; Zheng et al. 2021; Fumagalli et al. 2022). Notably, we found that virus-induced clinical disease occurred independently of control of lung viral burden, as mice lacking type I and type III interferon signaling exhibited increased viral replication in the lung without weight loss (Figure 4E-G). The discrepancy in virus-induced weight loss between interferon knockout mice and *Ly6e* knockout mice additionally suggests that constitutive *Ly6e* expression, which is present in interferon receptor knockout mice but not in *Ly6e* knockout mice, rather than interferon-induced *Ly6e*, which is absent in both interferon knockout mice and *Ly6e* knockout mice, may underly control of early viral pathogenesis in our SARS-CoV-2 infection model. From histopathological analyses we observed that *Ly6e* knockout mice exhibited a modest increase in lung pathology relative to wildtype mice, but this appeared to be driven by robust viral infection of the epithelium and not elevated recruitment of CD45+ immune cells (Figure 4). The striking increase in viral infection in *Ly6e* knockout lungs relative to wildtype controls by histopathology but not by plaque assay or quantitative PCR was likely due to differences in sample preparation: live virus in bronchiolar lavage fluid would not be detected in stained lung sections but would contribute to the overall viral burden quantified in homogenates made from whole, unflushed lungs. At the tissue level, *Ly6e* knockout mice exhibited a robust antiviral interferon response after SARS-CoV-2 infection that paralleled that of wildtype mice, but also expressed fewer secretory cell genes when compared to uninfected *Ly6e* knockout mice or infected wildtype mice (Figure 5, S4). SARS-CoV-2 has been previously shown to exhibit tropism for lower respiratory tract secretory club cells (Salahudeen et al. 2020; Ravindra et al. 2021; Fiege et al. 2021; Peng et al. 2022), which was associated with subsequent loss of the secretory club cell marker, *Scgb1a1*, in a mouse-adapted SARS-CoV-2 model (Leist et al. 2020). Infection of *Ly6e* knockout mice with a natural SARS-CoV-2 variant of concern may therefore be a useful tool for understanding how increased susceptibility of secretory cells results in clinical disease.

The significant reduction of *Muc5b* expression in SARS-CoV-2-infected *Ly6e* knockout mice relative to infected wildtype controls suggests that lower levels of this gene may contribute to the increase in pathology and virus-induced weight loss observed in knockout animals. In corroboration of this hypothesis, multiple meta-analyses of COVID-19 patient outcomes revealed a protective effect of the *MUC5B* polymorphism rs35705950-T, which increases gene expression, against severe disease and hospitalization (Fadista et al. 2021; van Moorsel et al. 2021; Verma et al. 2021). Thus, constitutive expression of *Ly6e* may protect respiratory tract secretory cells from SARS-CoV-2 infection and confer resistance to viral pathogenesis, possibly through preservation of *Muc5b* expression. While further mechanistic dissection is necessary to test whether reduced *Muc5b* expression mediates disease in *Ly6e*-knockout mice, Gene Ontology enrichment analysis highlights a possible effect on leukocyte chemotaxis and activation.

In aggregate, our study highlights compartment-specific antiviral defense in resistance to murine and human coronavirus infection. While our data demonstrates a conserved role for *Ly6e* in resistance to distinct coronaviruses, it also suggests an additional *Ly6e-* and interferon-independent defense mechanism that is highly protective from infection by SARS-CoV-2 but not murine coronavirus. Further investigation into the mechanism underlying natural murine resistance to SARS-CoV-2 may be useful for understanding the pleiotropic infection outcomes observed in human populations.

### Limitations of Study

Our approach of generating conditional knockout and bone marrow chimeric mice allowed us to identify distinct compartments that confer resistance to MHV infection through expression of *Ly6e*, such as *Lyz2*-expressing myeloid cells and epithelial cells, radioresistant *Vav1*-expressing cells, or non-hematopoietic cells in Sox2-Cre mice. However, we are unable to conclude whether *Ly6e* expression is required for host survival by a single cell type or multiple cell types from any group. Another major limitation of our study is that we do not provide direct evidence that secretory cells are targeted and killed by SARS-CoV-2 infection, which we infer from the significant reduction in secretory cell-associated genes in whole lung transcriptomes. Infection with mouse-adapted SARS-CoV-2 led to fewer bronchiolar cells expressing the traditional club cell marker *Scgb1a1* (Leist et al. 2020), but it is unclear if decreased expression of secretory cell-associated genes or proteins indicates virus-induced cell loss or transcriptional dysregulation.

## Materials and Methods

### Viruses

Generation of an infectious clone of mouse hepatitis virus, polytropic strain A59 (MHV-A59) has been described previously (Coley et al. 2005). MHV-A59 stocks used for infections were prepared by propagation of viral seed stocks in 17CL1 cells. In brief, a 100% confluent 175 cm^2^ tissue culture flask of 17Cl1 cells was inoculated with a concentrated stock of MHV-A59 at a MOI of 0.4 in 10 mL of 10% FBS (Gibco, MA)/MEM (Gibco, MA) for 3 to 5 hours at 33°C. Viral inoculum was replaced with 0.22 μm filter-sterilized pH 6.8 growth medium (1X non-essential amino acids [Gibco, MA]/20 mM HEPES, pH 6.8 [Gibco, MA]/5 mM sodium bicarbonate [Sigma-Aldrich, MA]/1X penicillin-streptomycin [Gibco, MA]/1% tryptose phosphate broth [Gibco, MA]/10% FBS [Gibco, MA]/DMEM #12100-046 [Gibco, MA]), to limit syncytial formation and cells were incubated overnight at 33°C then the entire flask was frozen at −20°C. After thawing the flask at room temperature, the supernatant was clarified by centrifugation at 1000 *xg* for 5 minutes before aliquoting and storage at −80°C. Viral titer was determined by plaque assay on L929 cells. In brief, L929 were plated on 24 well plates at a density of 165,000 cells per well the day before inoculation with 12 dilutions from a 10-fold dilution series. The infection was incubated for 2-3 hours at 37°C and then the viral inoculum was removed and replaced with a pH 7.2 methyl cellulose overlay medium (1% methyl cellulose [Sigma-Aldrich, MA]/5% FBS [Gibco, MA]/1X penicillin-streptomycin [Gibco, MA]/44 mM sodium bicarbonate [Sigma-Aldrich, MA]/DMEM #12100-046 [Gibco, MA]). After overnight incubation at 37°C, overlay medium was removed, cells were washed twice with distilled water, and stained for at least 0.5 hours with crystal violet (0.5% crystal violet [Sigma-Aldrich, MA]/20% methanol [Sigma, MA]) before removal and syncytium enumeration.

SARS-CoV-2, isolate hCoV-19/Japan/TY7-503/2021 (P.1) of an unknown titer was obtained (BEI Resources, VA) and inoculated onto Vero-E6-C1008 (ATCC, VA) plated at a density of 7 x 10^6^ cells in a 175 cm^2^ tissue culture flask in 2% FBS (Gibco, MA)/1X non-essential amino acids (Gibco, MA)/MEM (Gibco, MA) for 45 minutes at 37°C. Viral inoculum was removed and replaced with 2% FBS (Gibco, MA)/1X non-essential amino acids (Gibco, MA)/MEM (Gibco, MA) and cells were incubated for 3 days at 37°C before supernatants were clarified by centrifugation at 1000 *xg* for 5 minutes before aliquoting and storage at −80°C. Viral titer was determined by plaque assay on Vero-E6-C1008 cells (ATCC, VA). In brief, cells were plated at 650,000 cells per well of a 6 well plate and infected with 6 dilutions from a 10-fold dilution series in 1% FBS (Gibco, MA)/1X non-essential amino acids (Gibco, MA)/MEM (Gibco, MA) for 30 minutes at 37°C. Inoculum was then removed and replaced with Avicel overlay medium (5% FBS [Gibco, MA]/1X penicillin-streptomycin [Gibco, MA]/1X GlutaMAX [Gibco, MA]/1X Modified Eagle Medium, Temin’s Modification #11935-046 [Gibco, MA]/1.2% Avicel RC-591 [DuPont, DE]). Cells were incubated for 3 days at 37°C before overlay medium was removed and cells were fixed with 4% PFA (Sigma-Aldrich, MA) for 30 minutes at room temperature. Fixative was then removed, and cells were stained for at least 0.5 hours with crystal violet (0.2% crystal violet [Sigma-Aldrich, MA]/20% ethanol [Sigma-Aldrich, MA]) before removal and plaque enumeration.

### Mice

Generation of *Ly6e^fl/fl^* (#036173, The Jackson Laboratory, ME) and *Ly6e^ΔVav1^* mice was described previously (Pfaender et al. 2020). *Ly6e^fl/fl^* mice were crossed with LysM-Cre transgenic mice (a kind gift from Lora Hooper, UTSW) to generate *Ly6e^ΔLyz2^* mice. *Ly6e^fl/fl^* mice were crossed with CD11c-Cre transgenic mice (a kind gift from Lora Hooper, UTSW) to generate *Ly6e^ΔCD11c^* mice. *Ly6e^fl/fl^* mice were crossed with CD4-Cre transgenic mice (#022071, The Jackson Laboratory, ME) to generate *Ly6e^ΔCD4^* mice. *Ly6e^fl/fl^* mice were crossed with CD8α-Cre transgenic mice (#017562, The Jackson Laboratory, ME) to generate *Ly6e^ΔCD8a^* mice. *Ly6e^fl/fl^* mice were crossed with CD19-Cre transgenic mice (#006785, The Jackson Laboratory, ME) to generate *Ly6e^ΔCD19^* mice. *Ly6e^fl/fl^* mice (originally generated on *CD45^.2/.2^* background) were crossed with *CD45^.1/.1^* mice (#002014, The Jackson Laboratory, ME) to generate *Ly6e^fl/fl^; CD45^.1/.1^* mice. Whole body *Ly6e* knockout mice were generated as previously described (Langford et al. 2018). In brief, a male *Ly6e^fl/fl^* mouse was crossed to a Sox2-Cre female mouse (#008454, The Jackson Laboratory, ME) to generate *Ly6e^Δ/+^; Sox2-Cre+* mice. *Ly6e^Δ/+^; Sox2-Cre+* male mice were crossed to *Ly6e^fl/fl^* female mice to obtain *Ly6e^Δ/Δ^; Sox2-Cre+* mice at a 3:1 Mendelian ratio. *Ly6e^Δ/Δ^; Sox2-Cre+* male mice were also crossed to *Ly6e^fl/fl^* female mice to obtain *Ly6e^Δ/Δ^; Sox2-Cre+*mice at a 1:1 Mendelian ratio. *Ifnar*^-/-^ and *Ifnlr*^-/-^ mice on a C57BL/6J background were a kind gift from Julie Pfeiffer (UTSW) and Megan Baldridge (WUSTL). C57BL/6J mice (#000664, The Jackson Laboratory) were purchased and delivered at least one week before infection to allow for acclimation. Genotyping was outsourced to Transnetyx (TN). All Cre recombinase-expressing strains were maintained by crossing a Cre hemizygous mouse to a Cre noncarrier mouse to ensure the possibility of littermate controls. Every effort was made to design experiments that used age, sex, and litter-matched controls when available. Male and female mice were used at an equal ratio for all experiments when available between the age of 6 to 15 weeks of age. Animal studies were carried out in specific pathogen-free barrier facilities managed and maintained by the UTSW Animal Resource Center. MHV-A59 infections were performed in animal biosafety level 2 containment. SARS-CoV-2 infections were performed in animal biosafety level 3 containment. Facilities were maintained at an acceptable range of 68–79 °F at a humidity of 30–70% on a 12 hour dark/12 hour light cycle. All procedures used in this study complied with federal and institutional guidelines enforced by the UTSW Institutional Animal Care and Use Committee and the UTSW Institutional Biosafety Committee and were granted institutional approval after veterinary and committee review (protocols #2016-101828 and #2020-102987).

### Bone marrow chimeras

Five- to ten-week-old *Ly6e^fl/fl^; CD45^.2/.2^* and *Ly6e^ΔVav1^; CD45^.2/.2^* littermates were selected as bone marrow recipients. Six-to twelve-week-old *Ly6e^fl/fl^; CD45^.1/.1^* mice were used as sex-matched donors. Recipient *CD45^.2/.2^* mice were started on 0.4 mg/mL enrofloxacin drinking water 1 week before irradiation and continued treatment 3 weeks after irradiation. Recipient animals were subject to lethal total body irradiation with 900-950 cGy in a SARRP Small Animal Irradiator (Xstrahl, GA) in the UTSW Preclinical Radiation Core Facility and rested for 6 to 7 hours. During that time, femurs and tibias were harvested from euthanized donor mice and flushed with 1X PBS (Gibco, MA) to isolate bone marrow. Red blood cells were lysed by treatment with 1X RBC lysis buffer (Tonbo Biosciences, CA) and removed by centrifugation. Cell density was determined with Invitrogen Countess 3 Automated Cell Counter (Thermo Fisher Scientific, MA). Six hours after irradiation, recipient animals were lightly anesthetized with isoflurane and retro-orbitally injected with 2 to 5 million live donor bone marrow cells resuspended in 50 μL of 1X PBS (Gibco, MA). Donor animals were rested for a minimum of 6 weeks before MHV-A59 infection or up to 11 weeks for assessing *Ly6e* expression and degree of chimerism.

### In vivo infections, viral titering, and serum ALT

For intranasal (peripheral) MHV-A59 and SARS-CoV-2 infections, mice were weighed then intraperitoneally injected with an anesthetic cocktail of 80 mg/kg ketamine, 6 mg/kg xylazine, and 1X PBS. Anesthetized mice were inoculated intranasally with 30 μL of virus diluted in chilled 1X PBS (MHV) or administered undiluted in 2% FBS (Gibco, MA)/1X non-essential amino acids (Gibco, MA)/MEM (Gibco, MA) (SARS-CoV-2) via the left nostril and then placed on heat until ambulatory. For intraperitoneal (systemic) MHV-A59 infections, mice were weighed then intraperitoneally injected with 100 μL of virus diluted in 1x PBS. Infected mice were monitored daily for weight and mortality. Animals that lost more than 20% of their original body weight were euthanized per Institutional Animal Care and Use Committee guidelines.

For quantifying MHV-A59 viral load, whole lungs, an approximately 2 cm^3^ section of liver, whole spleen, and heart were homogenized in 600 μL 2% FBS (Gibco, MA)/MEM (Gibco, MA) and the right hemisphere of the brain, including cerebellum and olfactory bulb, was homogenized in 800 μL 2% FBS (Gibco, MA)/MEM (Gibco, MA) with a single 3.5 mm stainless steel UFO bead (NextAdvance, NY) in a Bullet Blender Lite (NextAdvance, NY). All tissues were weighed before homogenization. Tissue homogenates were clarified of debris by centrifugation at 4°C for 800 *xg* for 5 minutes, and aliquots for plaque assay were frozen at −80°C. Blood was collected by terminal cardiac puncture immediately after CO_2_ asphyxiation and transferred to serum gel blood tubes (Thermo Fisher Scientific, MA). After a minimum of 20 minutes at room temperature, coagulated blood was removed by centrifugation at 4,000 *xg* for 5 minutes, and serum aliquots for plaque assay were frozen at −80°C. Plaque assay for MHV samples was performed as described in the above section titled ‘*Viruses*’ with four dilutions from a 10-fold dilution series. ALT was measured in fresh, unfrozen serum using VITROS MicroSlide Technology by the UTSW Metabolic Phenotyping Core.

For quantifying SARS-CoV-2 viral load, whole lungs from infected mice were weighed then homogenized in 600 μL 2% FBS (Gibco, MA)/MEM (Gibco, MA) with 3.2 mm stainless steel beads (NextAdvance, NY) in a Bullet Blender Storm Pro (NextAdvance, NY). Lung homogenates were clarified of debris by centrifugation at 800 *xg* for 5 minutes then aliquots were frozen at −80°C neat for plaque assay or diluted at a 1 to 4 ratio in TRIzol (Sigma-Aldrich, MA) for RNA isolation and quantitative PCR. Plaque assay for SARS-CoV-2 samples was performed as described in the above section titled *‘Viruses.’* RNA was isolated using the Direct-zol RNA miniprep kit following manufacturer’s instructions (ZymoResearch, CA). A 20 μl reaction contained 5 μL RNA (1/10^th^ of elution), 5 μL TaqMan Fast Virus 1-Step Master Mix, and 1.8 μL SARS-CoV-2 or *Rpl32* primer/probe set containing 6.7 μM each primer/1.7 μM probe (final concentration of primer/probe were 600 nM/150 nM probe). SARS-CoV-2 primers and probe were designed as recommended by the Center for Disease Control (GA) and are as follows: Forward (SARS-CoV-2_N1-F), 5’ GACCCCAAAATCAGCGAAAT 3’/ Reverse (SARS-CoV-2_N1-R), 5’ TCTGGTTACTGCCAGTTGAATCTG 3’/ Probe (SARS-CoV-2_N1-P), 5’ FAM-ACCCCGCATTACGTTTGGTGGACC-BHQ1 3’. *Rpl32* primers were previously described (Brattelid et al. 2010) and are as follows: Forward, 5’ CACCAGTCAGACCGATATGTGAAAA 3’/ Reverse, 5’ TGTTGTCAATGCCTCTGGGTTT 3’/Probe, 5’ CCGCCAGTTTCGCTTAA 3’. All oligonucleotides for SARS-CoV-2 and positive control (*Rpl32*) gene amplification were synthesized by LGC Biosearch Technologies (CA). In vitro transcribed RNA was used to generate a standard curve for detection of SARS-CoV-2 from a 10-fold dilution series starting at 5 x 10^10^ copies of RNA. SARS-Cov-2 nucleocapsid (*N*) gene was amplified by PCR from a synthesized *N* gene fragment (IDT, IA) with primers (IDT, IA) that introduced a T7 promoter sequence on the 3’ end: Forward, 5’ TAATACGACTCACTATAGGGATGTCTGATAATGGACCCCAAAATCAGC 3’/Reverse, 5’ CTAATTGCGGCCGCTTAGGCCTGAGTTGAGTCAGCAC 3’. PCR product was purified using QIAquick PCR Purification Kit (Qiagen, MD). In vitro transcription was performed using T7 RiboMAX Express Large Scale RNA Production System following manufacturer’s protocol (Promega, WI). RNA was quantitated by nanodrop on DS-11 FX Spectrophotometer (DeNovix, DE). Reverse transcription was performed at 50°C for 5 minutes, followed by inactivation at 95°C for 2 minutes, and 40 cycles of PCR (95°C for 3 seconds, 60°C for 30 seconds) on a QuantStudio 3 (Applied Biosystems, MA).

### Ly6e expression in purified cells and whole tissue

To confirm loss of *Ly6e* expression in *Ly6e^ΔCD4^, Ly6e^ΔCD8a^*, and *Ly6e^ΔCD19^* conditional knockout mice relative to *Ly6e^fl/fl^* littermates, spleens from uninfected mice were pulverized on a 70 μM cell strainer with a plunger from a sterile 3 mL syringe. Cells were rinsed from the strainer with 1X penicillin-streptomycin (Gibco, MA)/RPMI (Gibco, MA) and pelleted at 450 *xg* for 4-5 minutes at 4°C. Red blood cells were lysed by treatment with 1X RBC lysis buffer (Tonbo Biosciences, CA) and removed by centrifugation. Cell density was determined with Invitrogen Countess 3 Automated Cell Counter (Thermo Fisher Scientific, MA). For sorting experiments, 20 x 10^6^ splenocytes were processed and stained in 15 mL conical tubes as follows: 30 minute incubation with 1:1000 Ghost Dye Violet 450 per manufacturer’s protocol (Tonbo Biosciences, CA), 10 minute incubation with 1:100 anti-CD16/CD32 (Tonbo Biosciences, CA), and 20 minute incubation with primary antibody cocktail (*Ly6e^ΔCD4^*: 1:800 anti-CD4-PECy5 [Tonbo Biosciences, CA]/1:400 anti-CD3ε-FITC [Tonbo Biosciences, CA]; *Ly6e^ΔCD8a^*: 1:200 anti-CD8α-PECy7 [Tonbo Biosciences, CA]/1:400 anti-CD3ε-FITC [Tonbo Biosciences, CA]; *Ly6e^ΔCD19^*: 1:800 anti-CD19-PECy5 [Tonbo Biosciences, CA]/1:400 anti-CD3ε-FITC [Tonbo Biosciences, CA]). Splenocytes were washed then sorted for viability and cell surface markers as shown in Figure S1A by the UTSW Immunology Flow Cytometry Core. After sorting, cells were pelleted and lysed for RNA isolation with RNAaqueous-Micro Total RNA Isolation Kit (Thermo Fisher Scientific, MA) per manufacturer’s protocol.

To confirm loss of *Ly6e* expression in *Ly6e^ΔLyz2^* and *Ly6e^ΔCD11c^* conditional knockout mice relative to *Ly6e^fl/fl^* littermates, bronchoalveolar lavage fluid was collected in 5 mM EDTA/1X PBS (Gibco, MA) as previously described via the trachea (Jhingran, Kasahara, and Hohl 2016) and plated on a poly-lysine-coated 12 well plate in PBS++ (Gibco, MA) in alveolar macrophage media (1X penicillin-streptomycin [Gibco, MA], 10% FBS [Gibco, MA]/0.1%β-mercaptoethanol [Sigma-Aldrich, MA]). After a 2 hour incubation at 37°C, wells were rinsed twice with warm PBS++ (Gibco, MA) to remove non-adherent cells. Cells were lysed 4 hours later for RNA isolation with RNAaqueous-Micro Total RNA Isolation Kit (Thermo Fisher Scientific, MA) per manufacturer’s protocol. Alveolar macrophage purity of >95% (based on positive staining for F4/80 and CD11c) was previously determined by flow cytometry during the optimization of this protocol.

To measure *Ly6e* expression in CD45+ and CD31+ lung cells in *Ly6e^ΔVav1^* and bone marrow chimeric mice relative to *Ly6e^fl/fl^* mice, whole lungs were excised, rinsed once in 1X PBS, and mechanically homogenized following manufacturer’s instructions using a gentleMACS Octo Dissociator (Miltenyi Biotec, Germany) with the Lung Dissociation Kit, mouse (Miltenyi Biotec, Germany). Lung homogenates were filtered through a 70 μM cell strainer, centrifuged at 450 *xg* for 5 minutes, then subject to treatment with 1X RBC lysis buffer (Tonbo Biosciences, CA). Cell density was determined with Invitrogen Countess 3 Automated Cell Counter (Thermo Fisher Scientific, MA). Up to 20 x 10^6^ lung cells were stained in 15 mL conical tubes as follows: 30 minute incubation with 1:1000 Ghost Dye Violet 450 per manufacturer’s protocol (Tonbo Biosciences, CA), 10 minute incubation with 1:100 anti-CD16/CD32 (Tonbo Biosciences, CA), and 20 minute incubation with primary antibody cocktail (CD45+ immune cells: 1:50 anti-CD45-VioGreen [Miltenyi Biotec, Germany]; CD31+ endothelial cells: 1:2000 anti-CD31-PE [BioLegend, CA]/1:50 anti-CD45-VioGreen [Miltenyi Biotec, Germany]/1:100 anti-EpCam-PECy7 [BioLegend, CA]). Lung cells were washed then sorted for viability and cell surface markers as shown in Figure S2A by the UTSW Immunology Flow Cytometry Core. After sorting, cells were pelleted and lysed for RNA isolation with RNAaqueous-Micro Total RNA Isolation Kit (Thermo Fisher Scientific, MA) per manufacturer’s protocol.

For *Ly6e* expression in spleen, brain, heart, lung, and liver, tissues were homogenized in 600 μL or 800 μL (brain only) TRIzol (Sigma-Aldrich, MA) with a single 3.5 mm stainless steel UFO bead (NextAdvance, NY) in a Bullet Blender Lite (NextAdvance, NY). Unclarified tissue homogenates were diluted fresh in TRIzol (Sigma-Aldrich, MA) at a 1 to 9 ratio, and frozen at −80°C. For RNA isolation, chloroform was added to thawed samples at a 1 to 5 ratio, shaken for 15 seconds, then centrifuged for 15 minutes at 12,000 *xg* at 4°C. The aqueous phase was transferred to a fresh tube, mixed with 1 equal volume of 70% ethanol, then transferred to a RNeasy Mini Kit filter column (Qiagen, MD). Samples were processed per manufacturer’s instructions with on column DNase I treatment (Qiagen, MD).

RNA was quantitated by nanodrop on DS-11 FX Spectrophotometer (DeNovix, DE). *Ly6e* expression (Forward, 5’ atcttcggggcctcttcac 3’/ Reverse, 5’ atgagaagcacatcagggaat 3’) was quantified relative to the housekeeping gene *Rpl32* (Forward, 5’ aagcgaaactggcggaaac 3’/ Reverse, 5’ taaccgatgttgggcatcag 3’) using one-step RT-qPCR with either QuantiFast SYBR Green RT-PCR Kit (Qiagen, MD) or QuantiNova SYBR Green RT-PCR Kit (Qiagen, MD) per manufacturer’s protocol on a QuantStudio 3 (Applied Biosystems, MA).

### Flow cytometric quantification of CD45.1 + and CD45.2+ cells in spleen, lung, and blood

Spleens from uninfected mice were pulverized on a 70 μM cell strainer with a plunger from a sterile 3 mL syringe. Cells were rinsed from the strainer with 1X penicillin-streptomycin (Gibco, MA)/RPMI (Gibco, MA), pelleted at 450 *xg* for 4-5 minutes at 4°C, then subjected to treatment with 1X RBC lysis buffer (Tonbo Scientific, CA). Whole lungs were excised, rinsed once in 1X PBS, and mechanically homogenized following manufacturer’s instructions using a gentleMACS Octo Dissociator (Miltenyi Biotec, Germany) with the Lung Dissociation Kit, mouse (Miltenyi Biotec, Germany). Lung homogenates were filtered through a 70 μM cell strainer, centrifuged at 450 *xg* for 5 minutes, then subject to treatment with 1X RBC lysis buffer (Tonbo Biosciences, CA). Blood was collected by cardiac puncture using syringes coated internally with 50 μL of 0.5 M EDTA (Gibco, MA) then subjected to three sequential treatments with 1X RBC lysis buffer (Tonbo Scientific, CA) to isolate mononuclear cells. Cell density was determined with Invitrogen Countess 3 Automated Cell Counter (Thermo Fisher Scientific, MA). About 500,000 cells were stained as follows: 30 minute incubation with 1:1000 Ghost Dye Violet 510 per manufacturer’s protocol (Tonbo Biosciences, CA), 10 minute incubation with 1:100 anti-CD16/CD32 (Tonbo Biosciences, CA), and 20 minute incubation with primary antibody cocktail (1:50 anti-CD45.1-PE, REAfinity [Miltenyi Biotec, Germany]/1:50 anti-CD45.2-VioBlue, REAfinity [Miltenyi Biotec, Germany]). Samples were analyzed using a S1000 Flow Cytometer with an A600 96-well plate high-throughput extension (Stratedigm, CA) and compensated using CellCapture v5 (Stratedigm, CA). Data were analyzed with FlowJo v9 (Tree Star, OR). The total population of CD45+ immune cells was obtained by gating both CD45.1+ and CD45.2+ populations together then separately gating for each congenic marker within the combined gate (Figure S2C)

### Histopathology

SARS-CoV-2 infected mice were euthanized by CO2 asphyxiation and lungs were inflated to full tidal volume with 10% neutral buffered formalin (Sigma-Aldrich, MA) via the trachea which was then clamped shut. Pleural cavity organs were removed and placed in 20 volumes 10% neutral buffered formalin and agitated on an orbital shaker for a minimum of 72 hours. Heart, thymus, trachea, and lymph nodes were removed, and formalin-fixed lungs were submitted in 70% ethanol to the UTSW Histo Pathology Core for paraffin embedding, sectioning, and staining with hematoxylin and eosin (H&E). Scoring of lungs from SARS-CoV-2 infected mice for acute pneumonia were graded on a 4-point system (0, not seen; 1, mild; 2, moderate; 3, severe) for the following criteria: perivascular inflammation, alveolar inflammation, pleural inflammation, hemorrhage, and edema, with a maximum possible score of 15. Pathological scoring was performed by an independent UTSW pathologist who was blinded to the experimental conditions. The scoring criteria were adapted from a published rubric (Dietert et al. 2017). Immunostaining was performed at HistoWiz Inc. (Brooklyn, NY) as follows. Immunohistochemistry (IHC) was performed on a Bond Rx autostainer (Leica Biosystems) with citric acid-based retrieval buffer with a pH of 6.0 and Heat Induced Epitope Retrieval (HIER) for 20 minutes. Lung sections were stained with primary antibodies to detect either SARS-CoV-2 nucleocapsid (GTX635686, GeneTex, CA) or CD45-positive cells (ab25836, Abcam, UK) then developed with DAB (3,3’-diaminobenzidine) which was followed by a hematoxylin counterstain. The slides were scanned and digitized by Leica AT2 scanner (Leica Biosystems, Wetzlar, Germany). The digitized whole slide images were analyzed by Halo software (V3.3, Indica Labs, Albuquerque, NM, USA) at Histowiz Inc. The Multiplex IHC module was used to calculate the number of positive cells per square millimeter of tissue and the H score. The histopathological features of target findings were analyzed and quantified. H score was generated following the convention below: weak positive pixels (1+) are highlighted in yellow, moderate positive pixels (2+) are highlighted in orange, and strong positive pixels (3+) are in red. The positivity threshold of weak, moderate, and strong staining was 0.42, 0.78, and 1.3 respectively in Halo. The minimum tissue OD is 0.037. The same segmentation threshold of positive staining in nucleus and cytoplasm of target proteins decided by visual inspection were utilized throughout the whole image sets to maintain consistency in evaluation. An example of the image analysis is shown in Figure S3B-C.

### RNA-Seq

Whole lungs were isolated from SARS-CoV-2 infected mice and homogenized in 600 μL TRIzol (Sigma-Aldrich, MA) with 3.2 mm stainless steel beads (NextAdvance, NY) in a Bullet Blender Storm Pro (NextAdvance, NY). Lung homogenates were clarified of debris by centrifugation at 800 *xg* for 5 minutes, diluted into fresh TRIzol (Sigma-Aldrich, MA) at a 1 to 9 ratio, and frozen at −80°C. For RNA isolation, chloroform was added to thawed samples at a 1 to 5 ratio, shaken for 15 seconds, then centrifuged for 15 minutes at 12,000 *xg* at 4°C. The aqueous phase was transferred to a fresh tube, mixed with 1 equal volume of 70% ethanol, then transferred to a RNeasy Mini Kit filter column (Qiagen, MD). Samples were processed per manufacturer’s instructions with on column DNase I treatment (Qiagen, MD). SARS-CoV-2 N gene levels were also quantified in these samples by qPCR as described in the above section titled ‘*In vivo infections, viral titering, and serum ALT*.’ Purified RNA was submitted to Novogene (CA) in DNA/RNA shield (Zymo Research, CA) for library preparation and RNA-Seq analysis. Messenger RNA was purified from total RNA using poly-T oligo-attached magnetic beads. After fragmentation, the first strand cDNA was synthesized using random hexamer primers, followed by the second strand cDNA synthesis using dTTP for generation of a non-directional library. The library was checked with Qubit and real-time PCR for quantification and bioanalyzer for size distribution detection. The clustering of the index-coded samples was performed according to the manufacturer’s instructions. After cluster generation, the library preparations were sequenced on an Illumina platform and paired-end reads were generated. Raw reads of fastq format were first processed through in-house perl scripts. In this step, clean reads were obtained by removing reads containing adapter and low-quality reads from raw data. Index of the reference genome was built and paired-end clean reads were aligned to the reference genome using Hisat2 v2.0.5. featureCounts v1.5.0-p3 was used to count the reads numbers mapped to each gene and FPKM of each gene was calculated based on the length of the gene and reads count mapped to this gene. Differential expression analysis was performed using the DESeq2 R package (1.20.0). The resulting P-values were adjusted using the Benjamini and Hochberg’s approach for controlling the false discovery rate. Gene Ontology (GO) enrichment analysis of differentially expressed genes was implemented by the clusterProfiler R package, in which gene length bias was corrected. GO terms with corrected P value less than 0.05 were considered significantly enriched by differential expressed genes. Volcano plots and heatmaps were prepared using GraphPad Prism version 9.1.0.

### Statistical analysis

Statistical analyses for all data except for RNA-Seq data were performed using GraphPad Prism version 9.1.0. Individual statistical tests are specified within the figure legends.

## Acknowledgements

We thank members of the Schoggins lab for useful discussions. We would also like to thank Mariano Aufiero and Tobias Hohl (MSKCC) for advice and feedback for bone marrow chimera studies, the lab of Lora Hooper (UTSW) for CD11c-Cre and LysM-Cre transgenic mice, the labs of Julie Pfeiffer (UTSW) and Megan Baldridge (WUSTL) for *Ifnar*^-/-^ and *Ifnlr*^-/-^ mice, respectively, the UTSW Animal Resource Center for training and animal husbandry, the UTSW Metabolic Phenotyping Core for analysis of serum samples for ALT levels and expertise, the UTSW Histo Pathology Core, and the UTSW Immunology Flow Cytometry Core.

---

**Supplementary Figure 1.**
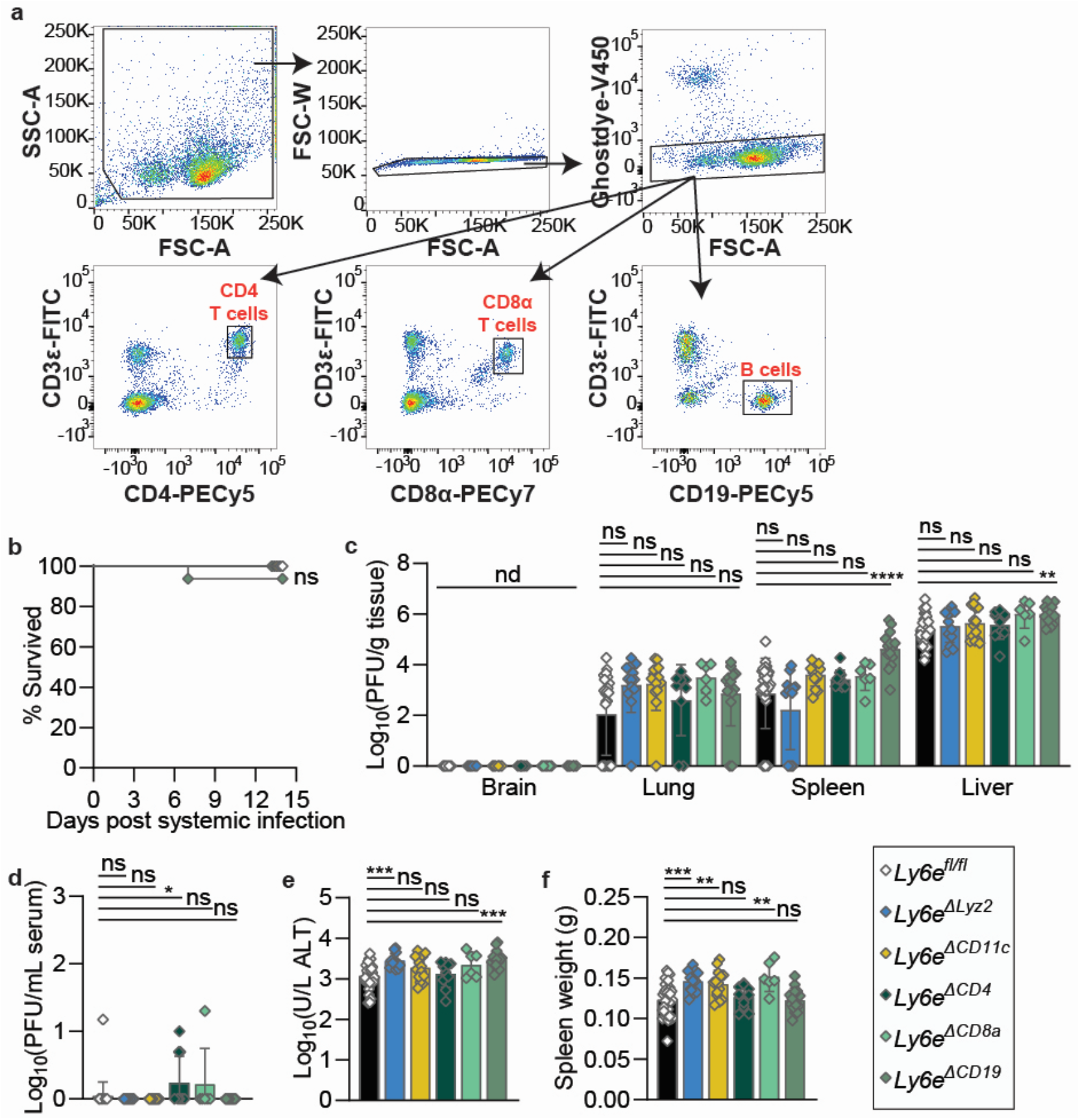
Cell-type specific contributions of Ly6e during systemic coronavirus infection. (A) Gating strategy for sorting lymphocytes from the spleen for examining Ly6e gene expression in Ly6e^ΔCD4^, Ly6e^ΔCD8a^, and Ly6e^ΔCD19^ mice relative to Ly6e^fl/fl^ littermates in Figure 2A. (B-F) Mice were systemically (intraperitoneally or IP) infected with 5,000 PFU MHV-A59 and assessed for (B) survival (n=47 Ly6e^fl/fl^, n=15 Ly6e^ΔLyz2^, n=14 Ly6e^ΔCD11c^, n=14 Ly6e^ΔCD4^, n=5 Ly6e^ΔCD8a^, and n=16 Ly6e^ΔCD19^), (C) viral burden in brain, lung, spleen, and liver, (D) viral burden in serum, (E) serum alanine aminotransferase, and (F) post-mortem spleen weight (n=30 Ly6e^fl/fl^, n=13 Ly6e^ΔLysM^, n=14 Ly6e^ΔCD11c^, n=10 Ly6e^ΔCD4^, n=6 Ly6e^ΔCD8a^, and n=14 Ly6e^ΔCD19^ for B-F). Male and female mice were used at an approximately 1 to 1 ratio for these experiments. Statistical significance was determined by log-rank (Mantel-Cox) tests (B), Kruskal-Wallis test (C-D), and one-way ANOVA (E-F). Error bars represent mean ± standard deviation. ns p>0.05, * p<0.05, ** p<0.01, *** p<0.001, **** p<0.0001.

**Supplementary Figure 2.**
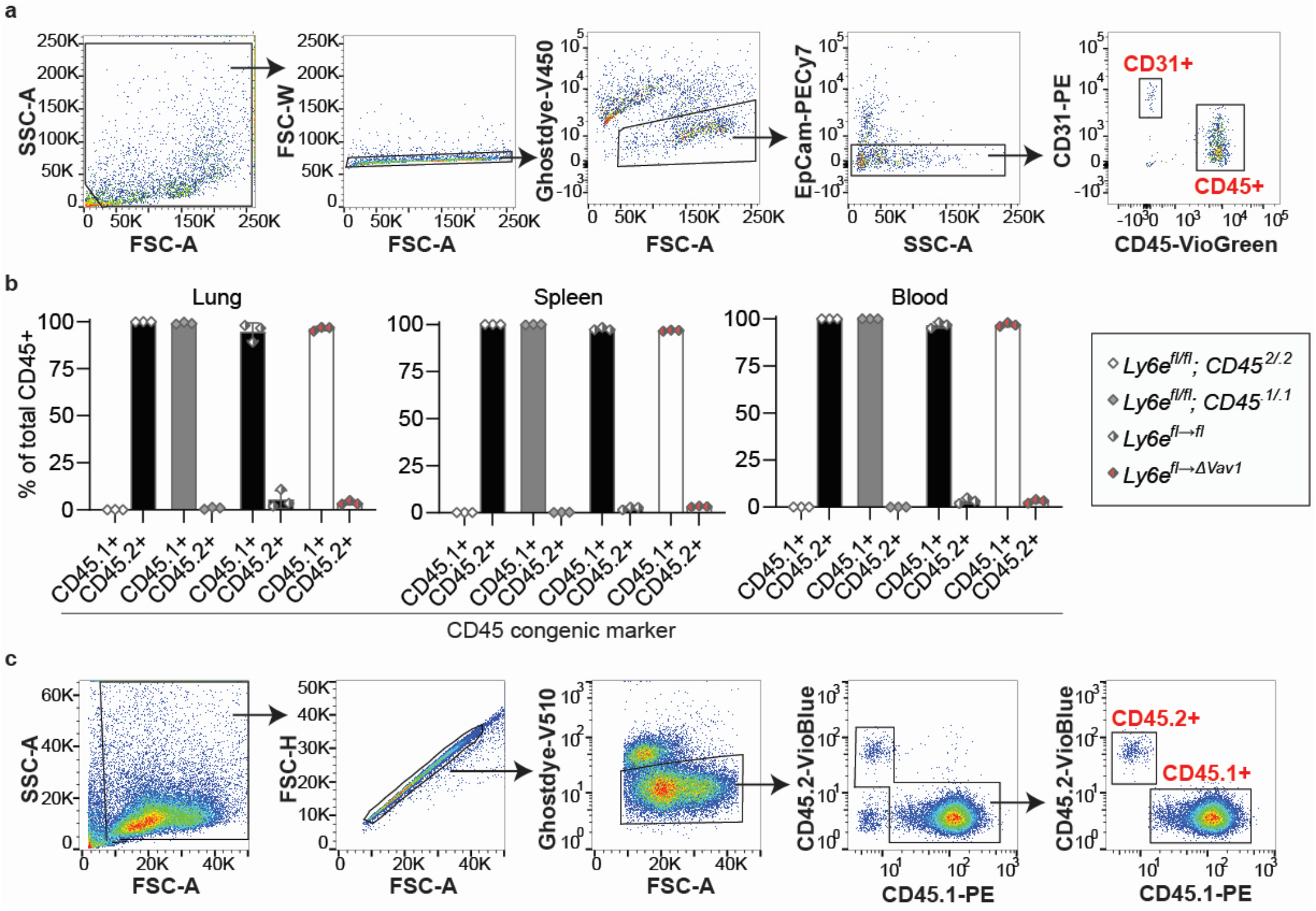
Gating strategy for Figure 3 and measuring composition of donor and recipient immune cells in bone marrow chimeric mice. (A) Gating strategy for sorting lung CD45+ cells and CD31+ cells from Ly6e^ΔVav1^ mice for determining Ly6e gene expression as shown in Figures 3C and 3D, respectively. (B) Relative composition of CD45.1+ and CD45.2+ immune cells of compartment total CD45+ cells in lung, spleen, and blood (n=3 Ly6e^fl/fl^; CD45^.2/.2^, n=3 Ly6e^fl/fl^; CD45^.1/.1^, n=3 Ly6e^fl→fl^, n=3 Ly6e^fl→ΔVav1^). (C) Gating strategy for (B).

**Supplementary Figure 3.**
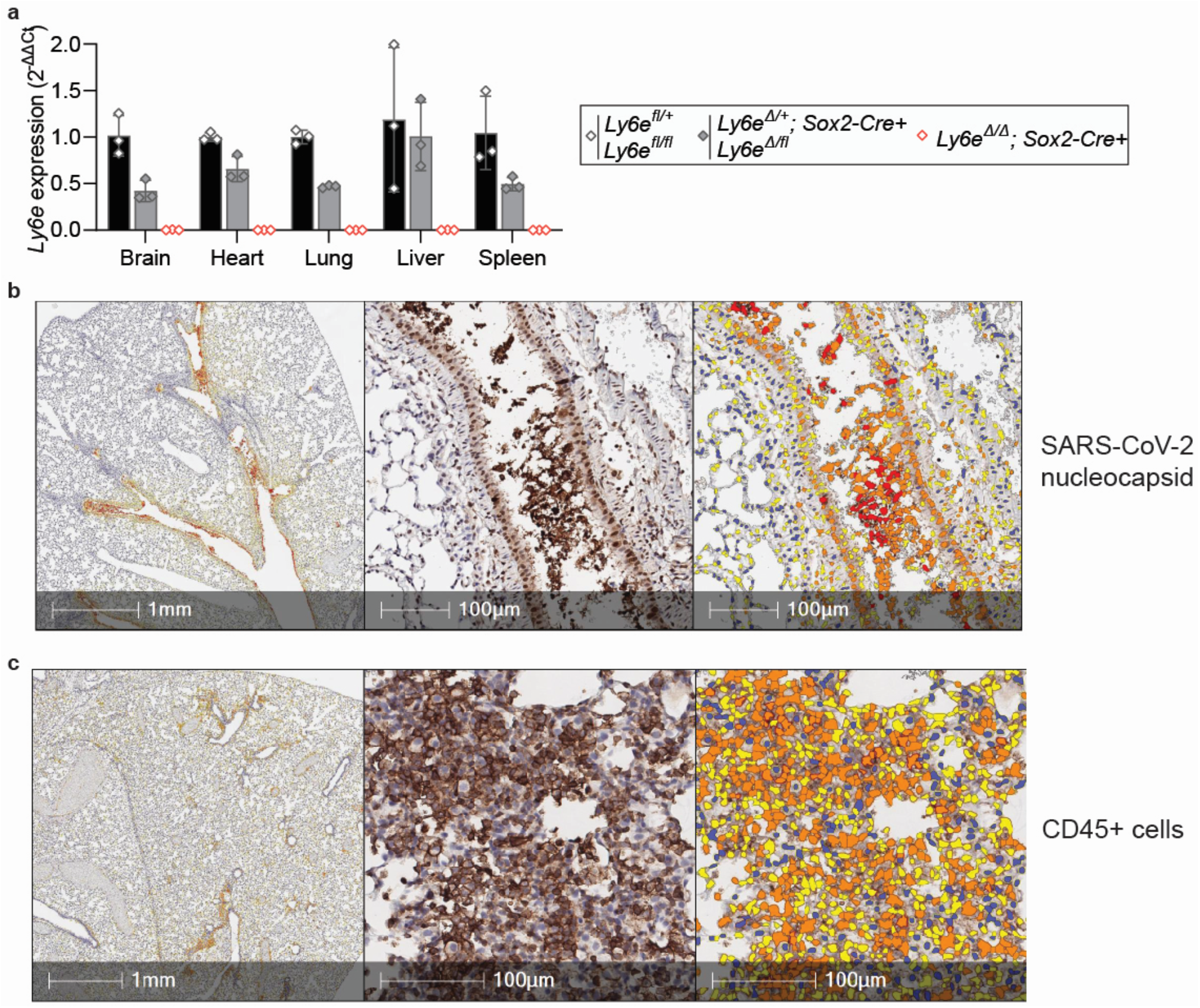
Ly6e gene expression in tissues and examples of image analyses for Figures 4K-N. (A) Relative Ly6e mRNA levels in brain, heart, lung, liver, and spleen (n=3 wildtype, n=3 heterozygous, and n=3 knockout). Example of automated image analysis of SARS-CoV-2 infected cells for Figures 4K-L (B) and of CD45+ cells for Figures 4M-N (C). In the corresponding markup images, cells without DAB marker (e.g. positive stain) are colored blue, and positive cells identified as weak positive, moderate positive, and strong positive for the DAB marker are colored as yellow, orange, and red respectively. Further details of the analysis are described in the Methods section titled ‘Histopathology.’

**Supplementary Figure 4.**
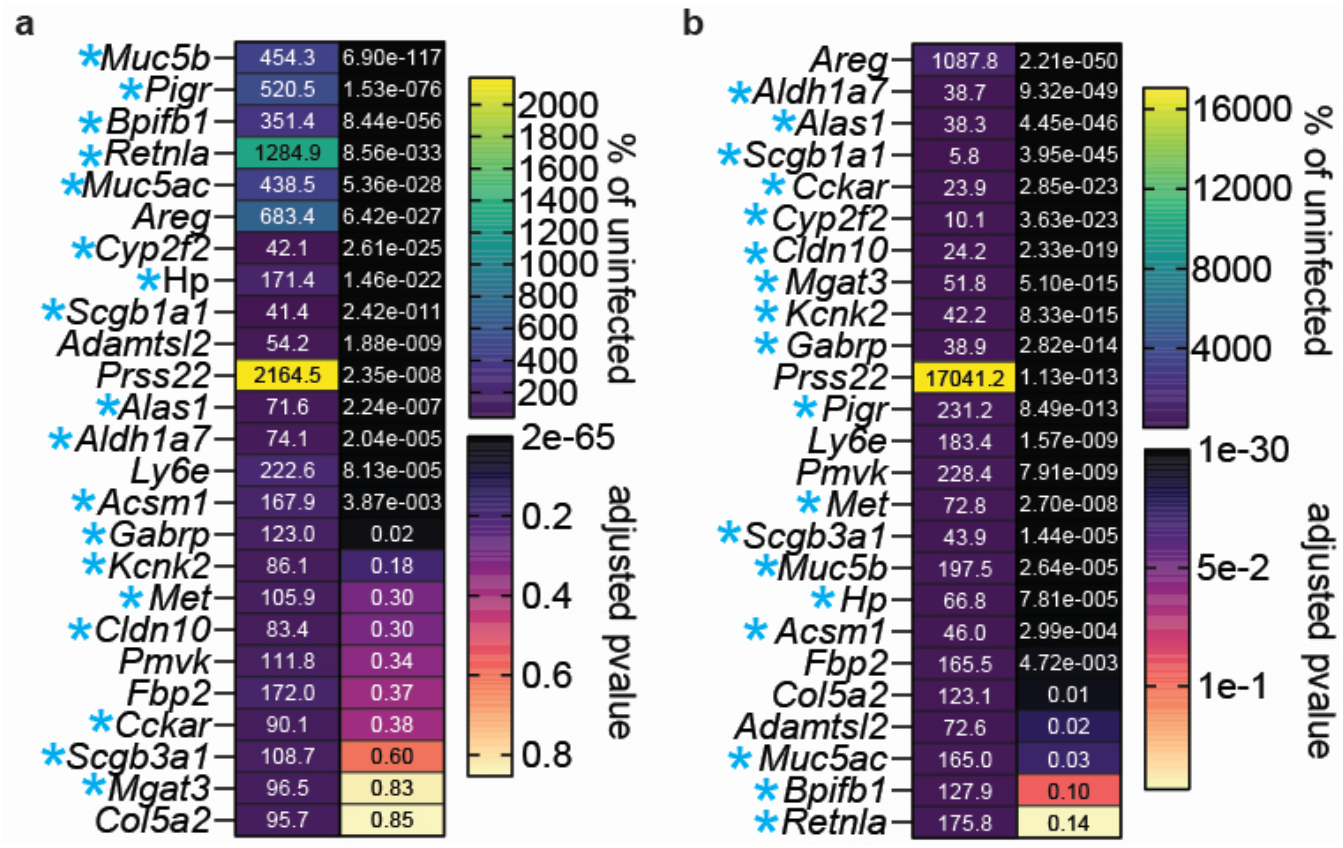
Expression of genes most differentially expressed between infected wildtype and knockout mice (Figure 4H) after SARS-CoV-2 infection. (A-B) Relative difference between SARS-CoV-2 infected and uninfected (A) wildtype mice and (B) knockout mice. Male and female mice were used at an approximately 1 to 1 ratio for these experiments. Genes expressed highly by secretory cells are indicated by a blue asterisk. Determination of statistical significance is described in the Methods section titled ‘RNA-Seq.’

**Supplementary Table 1.** Scoring for features of acute pneumonia for Figure 4.

| Mouse | Relative weight at 3 dpi | Perivascular inflammation (0-3) | Alveolar inflammation (0-3) | Pleural inflammation (0-3) | Hemorrhage (0-3) | Edema (0-3) | Total score (out of 15) |
| --- | --- | --- | --- | --- | --- | --- | --- |
| Wildtype 1 | 98.2 | 0 | 0 | 0 | 0 | 0 | 0 |
| Wildtype 2 | 100.0 | 1 | 0 | 0 | 0 | 0 | 1 |
| Wildtype 3 | 98.0 | 1 | 0 | 0 | 0 | 0 | 1 |
| Wildtype 4 | 92.3 | 1 | 0 | 0 | 0 | 0 | 1 |
| Wildtype 5 | 100.0 | 1 | 0 | 0 | 0 | 0 | 1 |
| Wildtype 6 | 101.4 | 1 | 0 | 0 | 0 | 0 | 1 |
| Wildtype 7 | 101.4 | 1 | 0 | 0 | 0 | 0 | 1 |
| Wildtype 8 | 103.8 | 1 | 0 | 0 | 0 | 0 | 1 |
| Wildtype 9 | 92.8 | 2 | 1 | 0 | 0 | 0 | 3 |
| Knockout 1 | 92.5 | 2 | 0 | 0 | 0 | 0 | 2 |
| Knockout 2 | 87.6 | 2 | 0 | 0 | 1 | 0 | 3 |
| Knockout 3 | 87.8 | 1 | 0 | 0 | 1 | 0 | 2 |
| Knockout 4 | 92.5 | 0 | 0 | 0 | 0 | 0 | 0 |
| Knockout 5 | 90.1 | 0 | 0 | 0 | 0 | 0 | 0 |
| Knockout 6 | 89.7 | 1 | 0 | 0 | 0 | 0 | 1 |
| Knockout 7 | 86.3 | 2 | 2 | 1 | 1 | 0 | 6 |
| Knockout 8 | 87.8 | 1 | 0 | 1 | 0 | 0 | 2 |
| Knockout 9 | 82.5 | 2 | 1 | 1 | 1 | 1 | 6 |

**Supplementary Table 2.** Automated quantitation of CD45 and SARS-CoV-2 nucleocapsid immunostaining for Figure 4.

| Mouse | Relative weight at 3 dpi | % CD45+ cells | CD45 H-score | % SARS-CoV-2 nucleocapsid+ | SARS-CoV-2 nucleocapsid H-score |
| --- | --- | --- | --- | --- | --- |
| Wildtype 1 | 98.2 | 27.1 | 32.7 | 0.1 | 0.1 |
| Wildtype 2 | 100.0 | 29.9 | 34.9 | 0.4 | 0.5 |
| Wildtype 3 | 98.0 | 28.1 | 33.0 | 0.6 | 0.9 |
| Wildtype 4 | 92.3 | 26.9 | 31.4 | 1.5 | 2.0 |
| Wildtype 5 | 100.0 | 27.2 | 31.6 | 0.5 | 0.6 |
| Wildtype 6 | 101.4 | 29.4 | 33.9 | 0.3 | 0.3 |
| Wildtype 7 | 101.4 | not stained | not stained | not stained | not stained |
| Wildtype 8 | 103.8 | not stained | not stained | not stained | not stained |
| Wildtype 9 | 92.8 | not stained | not stained | not stained | not stained |
| Knockout 1 | 92.5 | 26.0 | 31.3 | 11.7 | 15.6 |
| Knockout 2 | 87.6 | 32.0 | 42.1 | 15.4 | 20.1 |
| Knockout 3 | 87.8 | 27.6 | 34.1 | 7.6 | 10.6 |
| Knockout 4 | 92.5 | 30.2 | 36.9 | 5.4 | 6.6 |
| Knockout 5 | 90.1 | 21.2 | 24.6 | 6.9 | 8.6 |
| Knockout 6 | 89.7 | 26.8 | 31.5 | 8.2 | 10.0 |
| Knockout 7 | 86.3 | not stained | not stained | not stained | not stained |
| Knockout 8 | 87.8 | not stained | not stained | not stained | not stained |
| Knockout 9 | 82.5 | not stained | not stained | not stained | not stained |

